# Glassy Surfactants Enable Ultra-High Concentration Biologic Therapeutics

**DOI:** 10.1101/2024.09.09.612104

**Authors:** Carolyn K. Jons, Alexander N. Prossnitz, Noah Eckman, Changxin Dong, Ashley Utz, Eric A. Appel

## Abstract

Protein therapeutics, like peptides and antibodies, have become critical to healthcare. Despite their exceptional potency and specificity, biopharmaceuticals are prone to aggregation, often necessitating low formulation concentrations as well as cold storage and distribution to maintain stability. Yet, high doses are required to treat many diseases. To achieve these doses, most approved protein drug products are administered intravenously, imposing excessive burdens on patients and the healthcare system. New approaches are needed to formulate proteins at high concentrations to enable less burdensome subcutaneous injection, preferably in an autoinjector format. To address this challenge, we report a subcutaneously injectable biotherapeutic delivery platform composed of spray-dried protein microparticles suspended in a non-solvent liquid carrier. These microparticles contain only active biopharmaceutical agent and a high glass transition temperature polyacrylamide-derived copolymer excipient affording several key benefits over traditional excipients, including: (i) improved stabilization of biopharmaceuticals through the spray drying process, and (ii) improved morphology and properties of the spray-dried particles, enhancing suspension injectability. Experiments with albumin and antibodies demonstrate that this technology enables ultra-high-concentration protein formulations (exceeding 500 mg/mL) that are injectable through standard needles with clinically relevant injection forces. Additionally, experiments in mice with two clinically relevant antibody drugs show these ultra-high-concentration formulations reduce required injection volumes without altering pharmacokinetics or efficacy. This approach could nearly triple the number of commercial protein drugs amenable to subcutaneous administration, dramatically reducing burden and improving access to these critical biopharmaceuticals.

**One Sentence Summary:** Here we leverage a unique copolymer excipient to enable ultra-high concentration protein formulations with improved stability and amenable to subcutaneous injection that can reduce patient burden, lower costs, and improve access to critical drugs.

## 1. Introduction

Biopharmaceuticals are an impactful class of therapeutics used to treat a wide range of diseases including cancer, autoimmune disorders, and viral infections (*1, 2*). Monoclonal antibodies (mAbs) dominate the biopharmaceutical market comprising over 50% of the almost 200 biologic product approvals in the last four years and 80% of the global annual sales of protein-based drugs (*3*). The success of mAbs is attributable to their complex tertiary structure that enables highly selective targeting of key molecules improving both their efficacy and safety profiles (*4*). Moreover their long serum half-lives (typically around 21 days, but can reach to 70 days for some variants) prolongs the timeframe over which the drug remains within the therapeutic window (*5*). Unfortunately, the high molecular weight and delicate tertiary structure of mAbs requires them to be delivered parenterally, typically by intravenous (IV) or subcutaneous (SC) injection (*3, 6*). To avoid the burden and costs associated with IV infusion, many new mAb drug products have been developed which are administered by SC injection (*2, 7*). While SC injections are preferable on account of their simplicity and reduced burden, the volume of administration is limited (0.5-1.5 mL), requiring either formulation at high protein concentrations (>100 mg/mL) (*5, 8–10*) or coformulation with enzymes (e.g., hyaluronidase) that degrade the tissue sufficiently to enable higher dosing volumes (*11*). Furthermore, mAbs are amphiphilic and possess hydrophobic regions that are highly susceptible to irreversible denaturation and aggregation in solution (*5, 8, 12–14*), which are exacerbated at elevated concentrations even under benign storage conditions (*15, 16*). These challenges hinder the adoption and implementation of mAb therapeutics for treatment of many diseases, particularly in under resourced locations around the world (*17–19*).

Biologic drug products rely on excipients for adequate formulation stability. Surfactant excipients such as Tween 20 or 80 (also called polysorbates) are present in 94% of FDA approved high concentration mAb drug products as they can screen interactions with hydrophobic interfaces (e.g., air-water or water-plastic), thereby preventing protein denaturation and aggregation. We previously reported the development of a novel surfactant excipient comprising a polyacrylamide-derived copolymer poly(acryloylmorpholine-co-N-isopropylacrylamide) (MoNi) that exhibits dramatically improved stabilization of proteins, such as insulin and mAbs, during stressed aging (*20–24*). In addition to these surfactants, sugars, such as trehalose and sucrose, can act as hydrophilic co-solutes to crowd proteins and prevent aggregation (*25*). Lastly, buffering additives such as amino acids (e.g., histidine and arginine) have been shown to preserve protein structure and reduce formulation viscosity by minimizing electrostatic repulsion between mAbs (*26–28*). Unfortunately, the optimization and combination of these various classes of excipients has only enabled the commercialization of a few high concentration mAb drug products (100–200 mg/mL) for SC administration (*29*), demonstrating a critical need for alternative formulation approaches which simultaneously achieve sufficiently high protein concentrations (>350 mg/mL) and formulation stability to enable the majority of mAb drug products to be dosed SC.

To circumvent the challenges associated with standard aqueous formulations, an alternative approach using dried proteins suspended in non-solvents has recently been explored (*30*). It is well understood that drying antibodies, commonly with lyophilization in the presence of various excipients (e.g., trehalose), can stabilize the protein for years by preventing physical and chemical degradation (*31*). There are several clinically approved mAb products consisting of lyophilized protein and excipients that are reconstituted at high concentrations just before use (*2*). Unfortunately, reconstitution for these products can require upwards of 30 minutes, exposing the mAb to non-equilibrium conditions that induce aggregation (*5, 8, 9, 32, 33*). To avoid reconstitution, it is possible to either mill lyophilizate into injectable microparticles or directly dry protein and excipients into microparticles with spray drying processes (*30*). These protein microparticles are extremely stable and can form injectable suspensions when dispersed in a non-solvent (*34–36*). Yet even the spray dried suspension formulations reported to date have required at least 33 wt% of a high glass transition temperature (Tg) additive, such as trehalose, to stabilize the mAbs during the spray drying process and to achieve sufficiently glassy microparticles to form robust injectable suspensions (**SI Fig. 1**). As a result, these microparticle suspensions have typically required a high total solid content to match liquid counterparts (300 mg/mL solids for 200 mg/mL mAb). While dry protein suspensions hold significant promise for improving the accessibility and delivery of biotherapeutics, current formulations have failed to make a clinical impact due to the modest increases in protein concentration.

**Figure 1.**
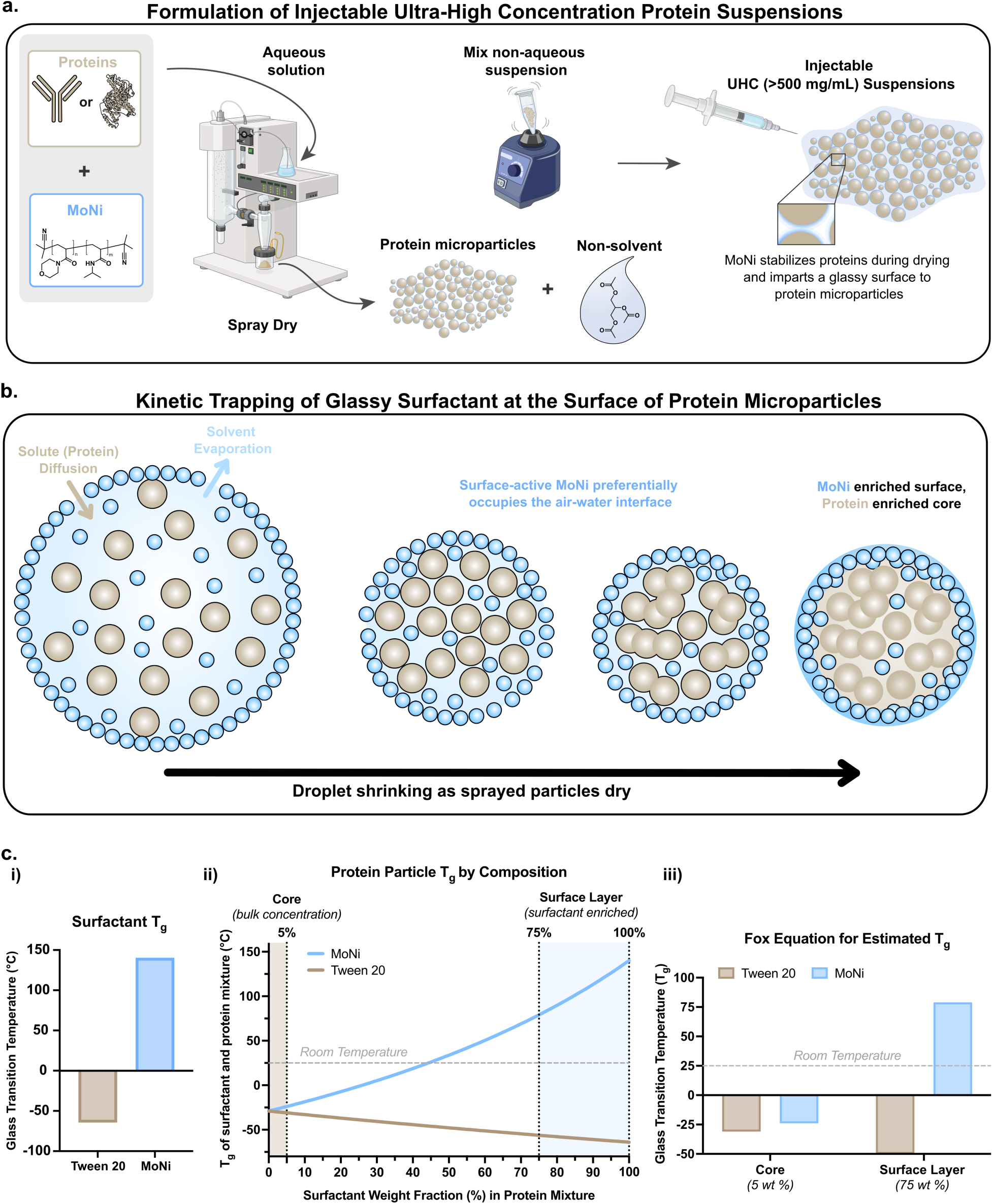
Overview of spray drying protein microparticles with MoNi excipient, the resulting distribution of components in the dried product, and estimation of glass transition temperatures of the microparticles. **a)** Schematic of spray drying process and formulation of ultra-high-concentration protein suspensions. **b)** Representation of surface-enrichment of MoNi surfactant that occurs as atomized droplets dry. Soluble protein diffuses towards the core and surface-active surfactants remain at the air-water interface. **c)** Estimation of glass transition temperature of protein microparticles as a function of composition at the surface and core of microparticles. **i)** Glass transition temperature for commercial surfactant Tween 20 compared to the polyacrylamide surfactant MoNi. **ii)** Estimation of the Tg of the protein surfactant mixture in the microparticles with the Fox equation. The core of the microparticles contains only 5% surfactant, while the surface can enrich up to 10-fold during the drying process (*37, 38, 42*). **iii)** The surface of microparticles stabilized with MoNi can easily achieve glass transition temperatures above room-temperature yielding hard glassy and injectable microparticles. Current commercial surfactants depress the glass transition temperature of the microparticles resulting in soft and tacky particle surfaces that stick to one another during injection.

We hypothesized that our surfactant excipient MoNi, which has an inherently high Tg on account of its polyacrylamide-derived chemical structure, could provide superior protein stability compared to standard sugar excipients at significantly lower concentrations in microparticle suspensions. Several studies have determined that surfactants are crucial additives for protein stability during spray drying since these molecules preferentially occupy air-water interfaces following aerosolization and exclude proteins from these hydrophobic interfaces more effectively than sugars and amino acids (*37–40*). Typical surfactants (e.g., Tween or Pluronics) are PEG-based and exhibit low Tg values (Tg ∼ -64 °C) that negatively impact the mechanical properties of the resulting microparticles (*41*). This generally requires the additional inclusion of high Tg sugars such as trehalose (Tg ∼ 106°C) to stabilize the particles at the expense of protein loading (*41*). In contrast, we hypothesize that a high Tg surfactant such as MoNi (*vide infra*) may encompass both the crucial functions of protecting proteins from unfavorable interface interactions during aerosolization and enhancing the mechanical properties of the microparticles to simultaneously improve suspension injectability and stability (**Fig. 1a**). In this work, we leverage the excipient MoNi to generate high protein content (∼95 wt%) spray-dried microparticles with both albumin and human IgG. We demonstrate that this approach enables ultra-high-concentration suspension formulations exceeding 500 mg/mL that are injectable through standard needles with clinically relevant injection forces. The MoNi excipient further stabilizes protein cargo during spray drying and storage as a liquid-suspension. Additionally, experiments in mice show these ultra-high-concentration formulations reduce required injection volumes without altering pharmacokinetics. Overall, we demonstrate that the high Tg excipient MoNi enables readily injectable protein formulations at the highest reported protein concentrations to date, nearly tripling the number of commercial protein drugs amenable to subcutaneous administration. These advancements could enable the development of drug products that dramatically reduce patient burden and improve access to critical biopharmaceuticals.

## 2. Results

In this work, we report the development of a biotherapeutic formulation comprising microparticles of protein and the MoNi excipient suspended in a non-solvent liquid carrier that is readily administered by injection through standard syringes and needles. We synthesized MoNi as previously reported and characterized the copolymer excipient with size exclusion chromatography (SEC) and ^1^H NMR (**SI Fig. 2**) (*23*). Additionally, differential scanning calorimetry (DSC) measurements demonstrated that MoNi exhibits a Tg of 140 °C (**SI Fig. 2**). For biotherapeutic formulation, protein and excipient can be directly dried from an aqueous solution into microparticles through spray drying (**Fig. 1a**) (*43*). In these processes, an aqueous feed solution is atomized to form small droplets, which are then exposed to hot air within the spray drying chamber leading to rapid liquid evaporation, and the resultant dried microparticles are separated and collected using a cyclone separator (*44, 45*). As only volatile compounds such as the aqueous media are driven off during the drying process, the bulk composition of the final microparticles is determined by the ratio of protein and excipients added to the initial aqueous feed solution, and the microparticle surface composition is determined by the relative surface-activity of protein and excipients (e.g. trehalose, surfactants) (*37, 38, 46*).

**Figure 2.**
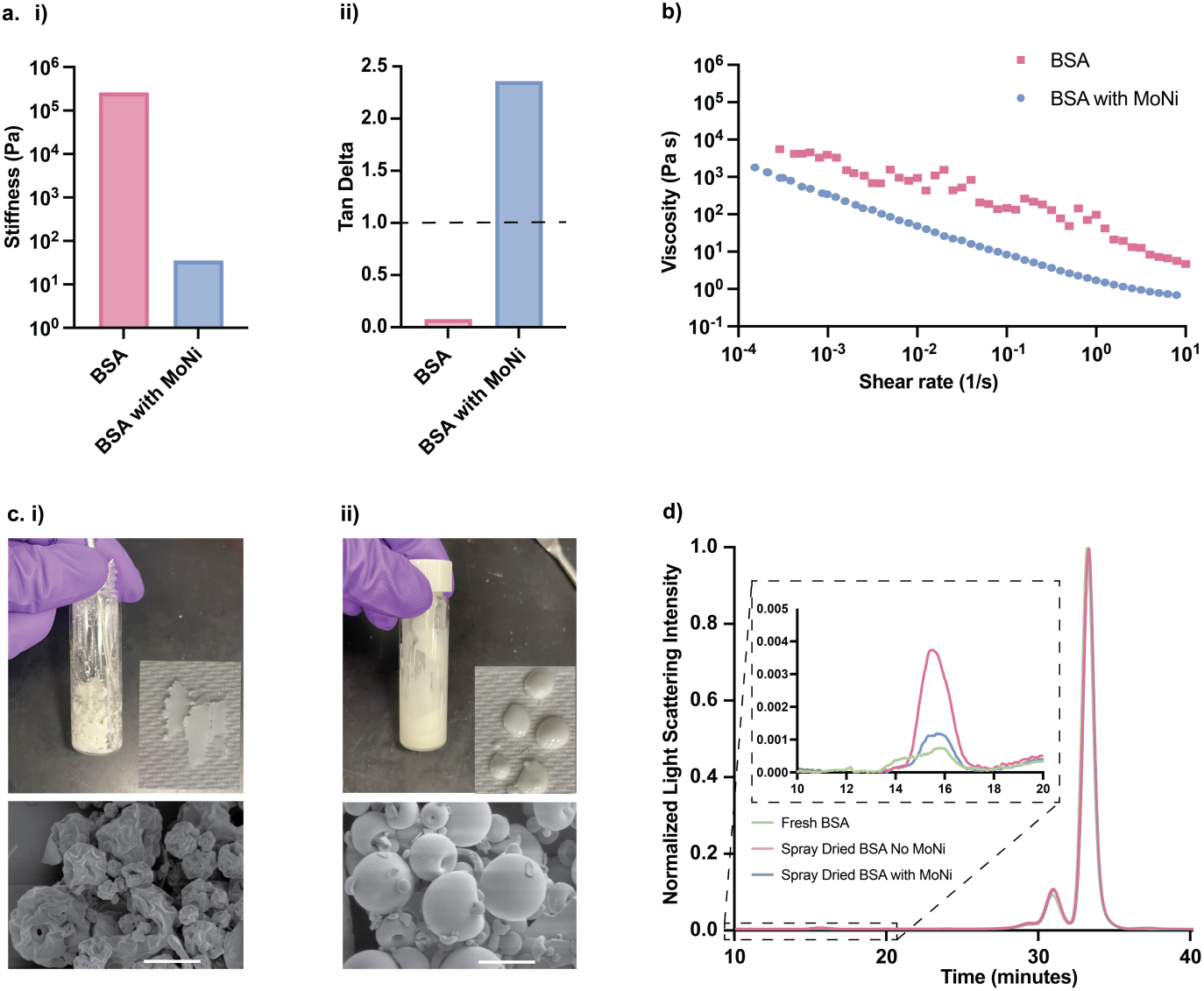
Rheological and stability characterization of spray dried microparticles. **a)** Comparative **i)** storage modulus and **ii)** tan delta of suspensions at 10 rad/sec. **b)** Flow sweep of 520 mg/mL BSA microparticles resuspended in triacetin and 520 mg/mL BSA microparticles containing 5 wt% MoNi resuspended in triacetin. **c)** SEM image of microparticles (scale bar 5 μm) and image of resultant suspension in vial and on a glass slide in triacetin at 520 mg/mL for microparticles formulated **i)** without and **ii)** with 5 wt% MoNi. **d)** SEC trace of fresh BSA control, spray dried BSA without MoNi, and spray dried BSA with 5 wt% MoNi. PBS with sodium azide is used as an eluent.

Here, we spray dried aqueous solutions of either bovine serum albumin (BSA) or human IgG (hIgG) with MoNi excipient at a feed ratio of 20:1 (w/w) resulting in microparticles that are approximately 95 wt% protein and 5 wt% MoNi. While the resulting microparticles are only 5 wt% MoNi by bulk, the surface composition of the microparticles is expected to be heavily enriched with MoNi on account of its high surface activity (**Fig. 1b**). When considering a 10 μm protein microparticle and defining a surface depth of ∼5 nm, only 0.3 wt% loading of a highly surface-active additive would be needed to completely occupy the surface layer. Thus, the MoNi surfactant loading in our system is in roughly 15-fold excess the amount necessary to dominate the surface composition. At a concentration of 5 wt%, the properties of the microparticle surface will be dominated by the Tg of the surfactant (**Fig. 1c)**. Indeed, a conservative estimate of the surface properties of microparticles prepared with either protein alone, protein with MoNi, or protein with a standard PEG-based surfactant Tween 20 indicates that the MoNi is expected to yield robust, glassy particles, while particles formed from either protein alone or with Tween 20 would be soft and tacky (**Fig. 1ciii)**.

When spray dried particles prepared with BSA and MoNi were dispersed in a non-solvent liquid carrier such as triacetin at a concentration of 520 mg/mL BSA via vortexing, a stable fluid suspension was formed (**Fig. 2**). In contrast, when BSA particles prepared without MoNi were dispersed in triacetin at the same BSA concentration, a thick and solid paste was formed (**Fig. 2**). To ensure accurate protein concentrations in suspension formulations, the density of the BSA microparticles was determined by pycnometer (*vide infra*). Triacetin was chosen as a non-solvent in these initial experiments on account of its high density (1.16 g/mL), low viscosity (23 cP), low vapor pressure (0.0024 mm Hg), and inclusion in FDA-approved parenteral drug products (*47, 48*). Rheological characterization of the BSA microparticle suspensions in triacetin demonstrated that the inclusion of MoNi in the spray-dried particles significantly improves the flow properties of the formulation. Angular frequency sweeps demonstrated that the inclusion of MoNi in the BSA particles reduced the storage modulus values of the resulting suspensions by 4 orders of magnitude. Moreover, the Tan Delta (G’’/G’) values of the suspension formulation comprising MoNi exceeded 1, indicating liquid-like behavior, while the values for the suspension without MoNi were well below 1, indicating solid-like behavior (**Fig. 2a, SI Fig. 3**). Steady shear flow sweeps further demonstrated that the BSA suspensions are shear thinning, and that formulations comprising MoNi exhibited both lower viscosities and smoother flow than comparable formulations without MoNi (**Fig. 2b**). SEM images of spray dried microparticles illustrate that while BSA microparticles spray dried with and without MoNi were spherical and similar in average diameter (5-10 μm), inclusion of MoNi induced a smoother surface morphology (**Fig. 2c**). Beyond improving the physical characteristics of the BSA suspensions, inclusion of MoNi also stabilized the BSA protein through the spray drying process. Indeed, SEC characterization of fresh BSA as well as BSA spray dried with and without MoNi revealed that spray drying with MoNi resulted in both a higher monomer peak fraction and negligible formation of high molecular weight aggregates (**Fig. 2d**). These improvements in BSA stability and suspension flow behaviors with inclusion of MoNi in the particles were not unique to spray dried microparticles but were also observed when protein microparticles were instead formed by lyophilization followed by ball milling **(SI Fig. 4)**.

**Figure 3.**
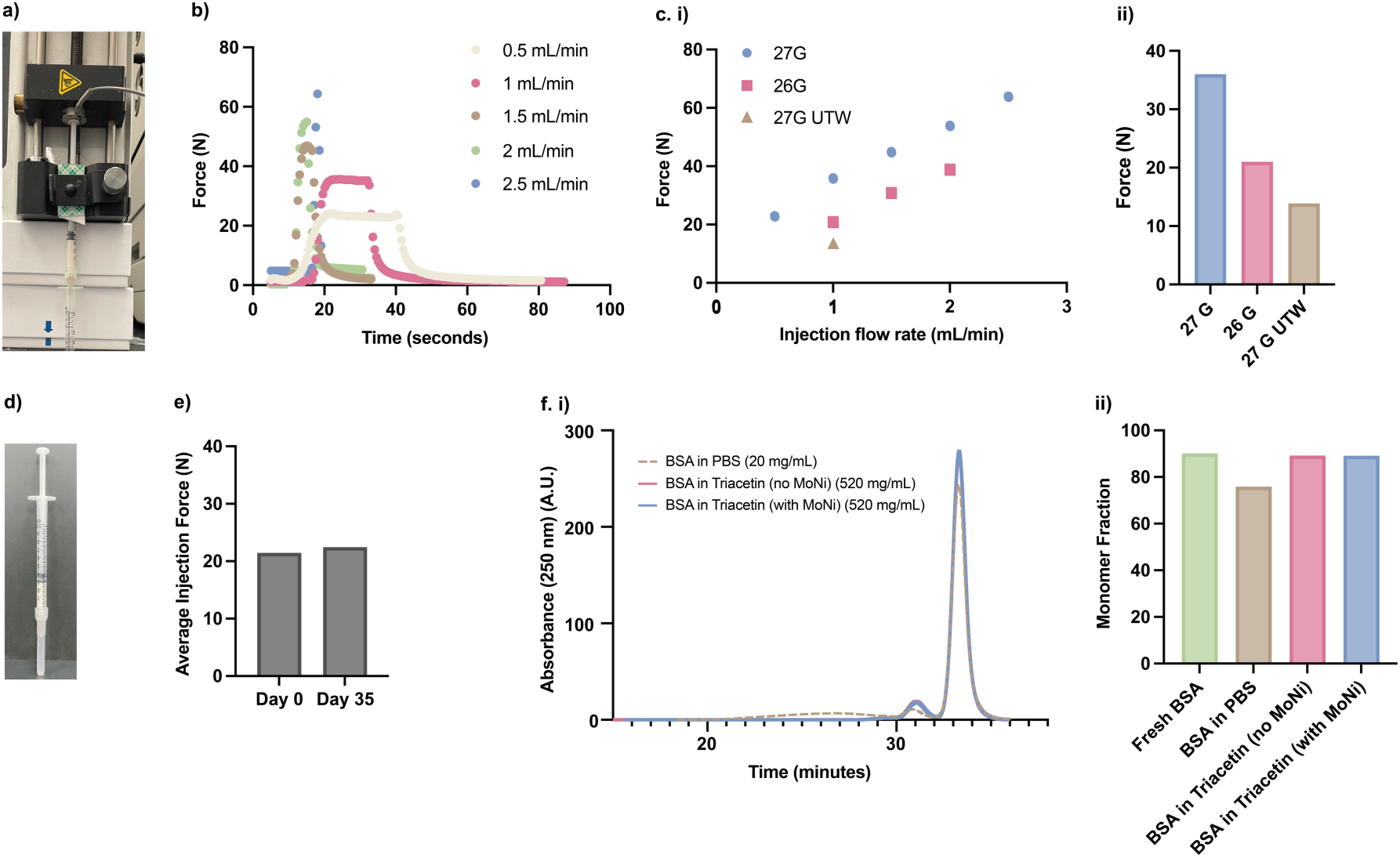
High-concentration protein formulations in triacetin are injectable and stable. **a)** Image of injection force setup (load cell connected to a syringe pump) used to quantify the force of injection using a 1 mL syringe through standard needle gauges. **b)** Injection force curves for injection of 520 mg/mL BSA, 5 wt% MoNi suspensions through 27G ½ inch needles at varied flow rates. **c. i)** Force of injection is linear with flow rate and **ii)** scales with needle gauge. **d)** Illustration of 520 mg/mL BSA with 5 wt% MoNi in triacetin following vertical storage in a syringe for 120 hours. **e)** Injection force of 520 mg/mL BSA with 5 wt% MoNi in triacetin at day 0 and day 35 using a 26G ½ inch needle and a flow rate of 1 mL/min. **f. i)** SEC trace of BSA at 20 mg/mL in PBS, spray dried BSA without MoNi at 520 mg/mL in triacetin, and spray dried BSA with 5 wt% MoNi at 520 mg/mL in triacetin following stressed aging for 30 minutes at 60 °C and **ii)** corresponding monomer fraction for each formulation following stressed aging.

**Figure 4.**
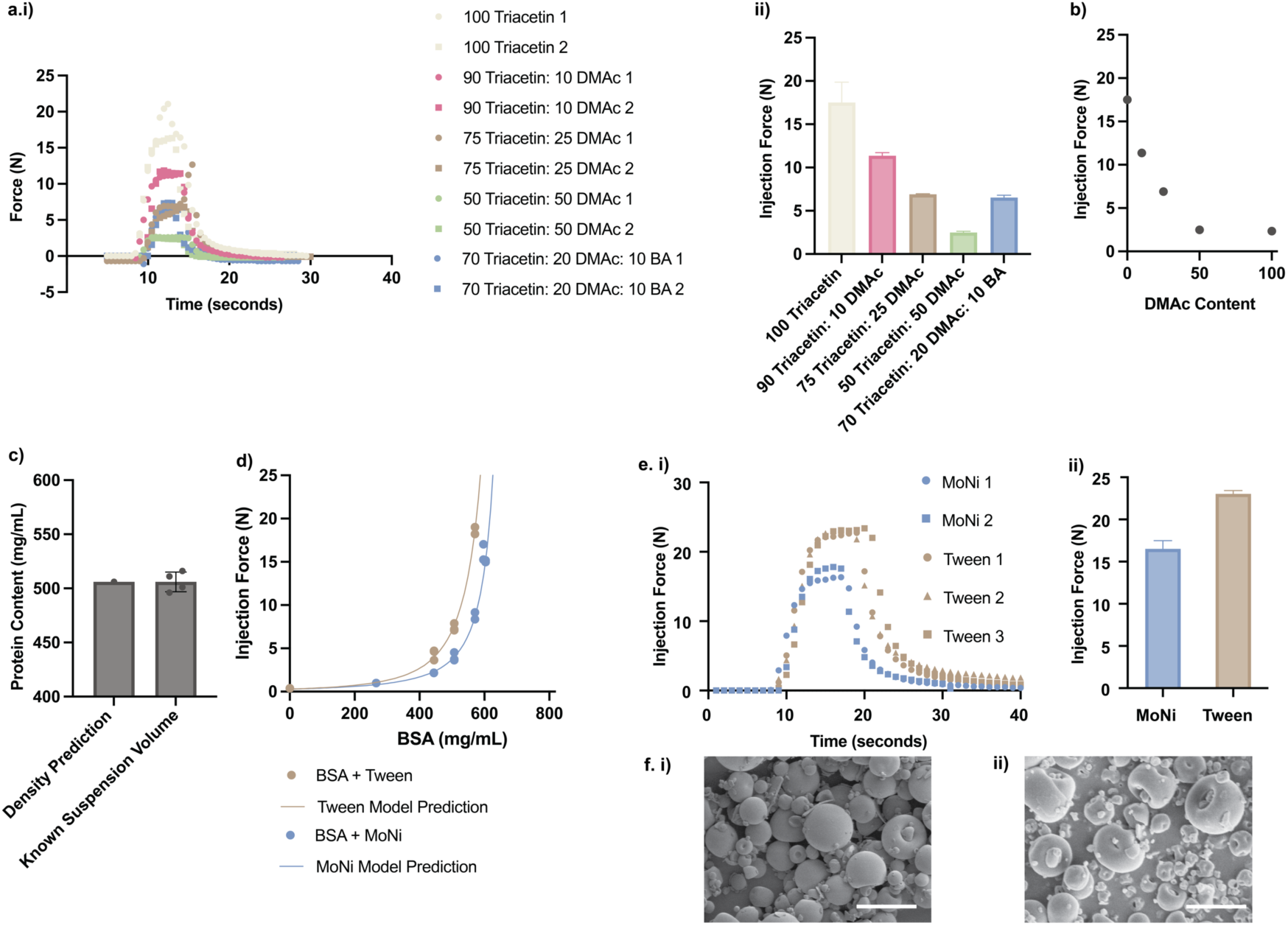
Low viscosity non-solvent additives allow for ultra-high-concentration protein formulations. **a. i)** Injection force curves for injection of 506 mg/mL BSA, 5 wt% MoNi suspensions at 1 mL/min through 26G ½ inch needles with DMAc and BA non-solvent additives, and **ii)** corresponding plateau injection forces (n=2). **b)** Plateau injection force of 506 mg/mL BSA, 5 wt% MoNi suspensions at 1 mL/min through a 26G, ½ inch needle as a function of DMAc content in triacetin (n=2). **c)** Comparative protein concentration as predicted from density measurements and measured via absorbance with nanodrop (n=4) from a known volume of suspension. **d)** Injection force as a function of concentration for BSA microparticles (with mol% matched MoNi or Tween 80) in 70 Triacetin: 30 DMAc. Injection force as a function of particle concentration is fit to a particle jamming model. **e. i)** Comparative injection force curves and **ii)** plateau injection forces for 506 mg/mL BSA microparticle suspensions (with mol% matched MoNi or Tween 80) in triacetin at 1 mL/min through 26G ½ inch needles (n=2). **f.** SEM of particle morphology for **i)** BSA MoNi and **ii)** BSA Tween microparticles (scale bars are 10 μm). Bar graphs show mean ± SD.

For therapeutic applications in SC injection, biopharmaceutical drug products need to be injectable through clinically relevant needle gauges with clinically relevant injection forces (*11, 34, 36*). To assess the injectability of the fluid-like suspensions, the injection force required to inject a suspension of BSA-MoNi microparticles in triacetin at a BSA concentration of 520 mg/mL through a 27 gauge, ½ inch needle at relevant flow rates was quantified using a force sensor attached to a syringe pump (**Fig. 3a**). In these assays, we observed that suspension injection force increases linearly with flow rate of injection, and decreases with the use of either smaller gauge needles or thin-walled needles, both of which have larger inner needle diameters (**Fig. 3b,c**). Using a clinically relevant ultrathin-walled (UTW) 27 gauge needle and a flow rate of 1 mL/min, the 520 mg/mL BSA suspensions comprising MoNi exhibited an injection force of only 14 N (**Fig. 3c**) – well within the injection force range achievable in standard pen autoinjectors (typically 25-30 N maximum force) (*49*). Lastly, long-term storage experiments to assess the physical stability of the suspensions demonstrated that they exhibit minimal particle settling in the triacetin non-solvent (**Fig. 3d**), and comparable injection forces (21.4 N vs. 22.4 N) over the course of 35 days of storage at room temperature (25 °C) (**Fig. 3e**).

In addition to the physical stability of the suspensions, the stability of the BSA protein itself in the triacetin-based suspensions through stressed aging was assessed. In these studies, BSA microparticle suspensions were prepared at 520 mg/mL protein in triacetin using spray dried BSA particles either with or without MoNi. An aqueous control solution was prepared with BSA at 20 mg/mL in PBS. The three formulations were heated at 60 °C for 30 minutes, and the BSA stability was then assessed by SEC (**Fig. 3f i**). While standard aqueous formulations of BSA in PBS showed significant aggregation and loss of monomer peak fraction (only 76% remaining), the BSA microparticle suspensions dispersed in non-solvent remain exceptionally stable with monomer peak fractions exceeding 89% through the stressed aging (c.f. 90% for fresh BSA; **Fig. 3f ii**). To further evaluate the stability of BSA in the triacetin-based microparticle suspensions prepared with and without MoNi, suspensions were stored at 4, 25, and 37 °C and BSA stability was assessed by SEC over time. Suspensions comprising MoNi showed smaller high molecular weight aggregate peaks than suspensions of BSA alone, indicating that the MoNi excipient improves the protein’s stability even in the dry state within the microparticles (**SI Fig. 5**).

**Figure 5.**
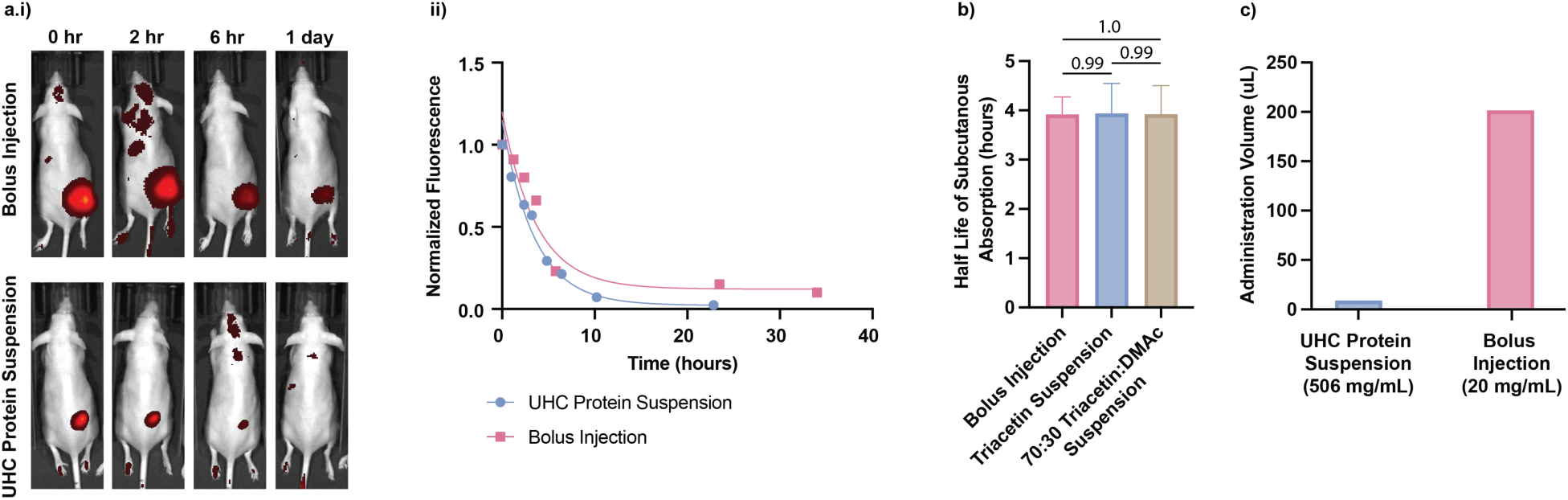
UHC BSA suspensions allow for in vivo administration and subcutaneous absorption. **a. i)** Representative IVIS images demonstrating subcutaneous absorption of fluorescently-tagged BSA administered via a PBS bolus injection or a UHC protein suspension. **ii)** Fluorescent signal in the subcutaneous space is fit to a single phase exponential decay mode to identify half-life of subcutaneous absorption. **b)** Comparative half-life of subcutaneous absorption for BSA administered via PBS bolus injection or UHC protein suspensions formulated with triacetin or 70 Triacetin: 30 DMAc (n = 3-5). **c)** Comparative volume of administration for bolus injection and UHC protein suspension. Bar graph shows mean ± SEM, statistical comparison using GLM followed by Tukey HSD.

To further reduce the injection force of protein suspensions and enable ultra-high-concentration (UHC) protein formulations, we evaluated non-solvent mixtures comprising triacetin and lower viscosity additives including dimethylacetamide (DMAc) and benzyl alcohol (BA) (*50, 51*). The notation “X Triacetin: Y DMAc: Z BA’’ indicates the volume fraction of Triacetin, DMAc, and BA in the non-solvent mixtures evaluated (**Fig. 4a i,ii**). For BSA-MoNi suspensions prepared at an equivalent protein content, the injection force at a given flow rate was reduced from 17.5 N to 2.5 N (7-fold reduction in injection force) with the use of alternative non-solvent additives (**Fig. 4a,b**). We selected 70 Triacetin: 30 DMAc (V/V) as our preferred non-solvent mixture as it afforded a greater than 2-fold reduction of injection force while demonstrating minimal SC injection site irritation in mice (**SI Fig. 6**).

**Figure 6.**
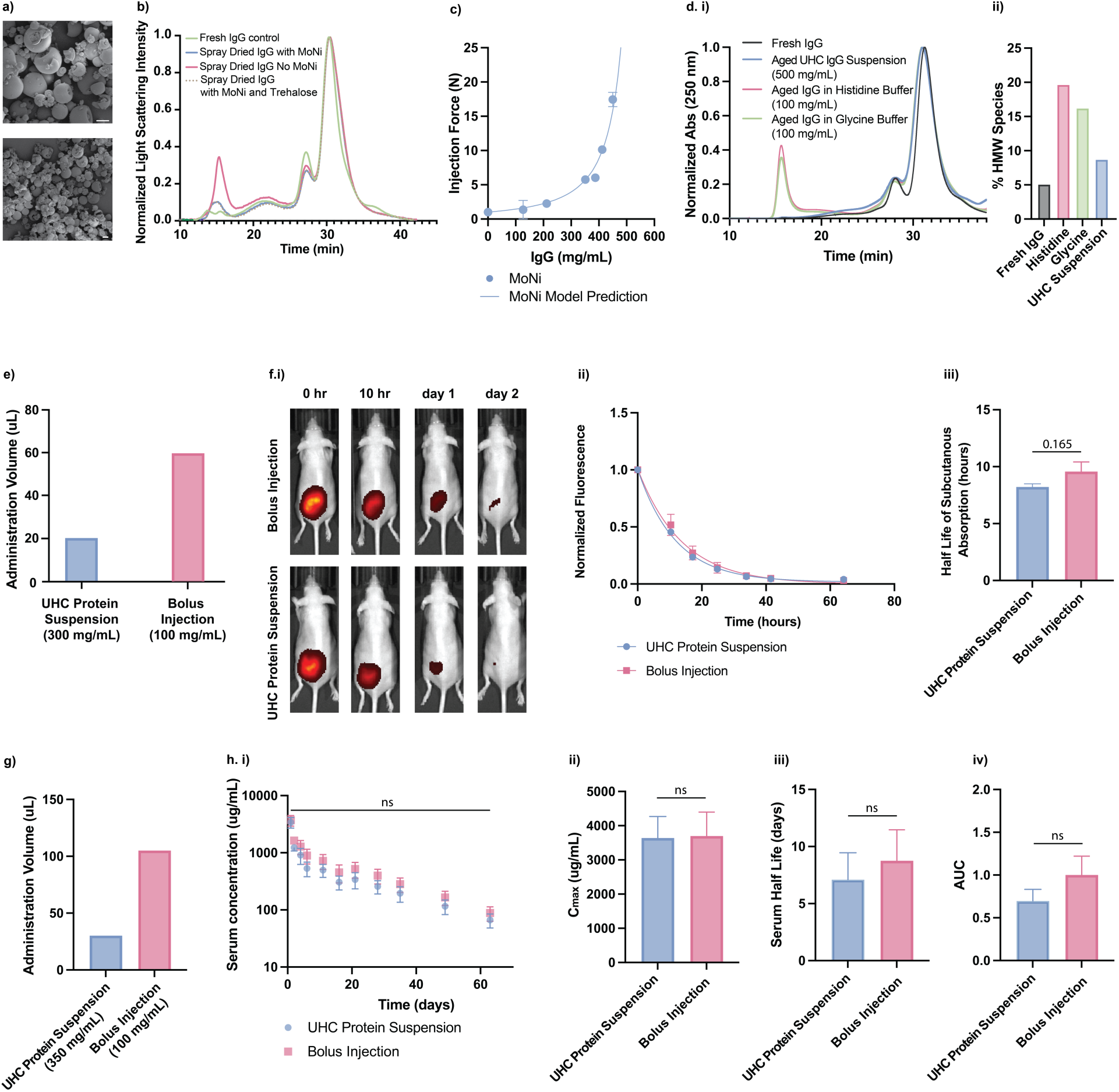
UHC suspension technology can be effectively used to deliver hIgG. **a)** SEM of spray dried IgG with MoNi particle morphology (scale bars are 10 μm). **b)** SEC trace of fresh hIgG control, spray dried hIgG without MoNi, spray dried hIgG with 5 wt% MoNi, spray dried higG with 25 wt% trehalose and 5 wt% MoNi. **c)** Injection force as a function of concentration for hIgG microparticles with MoNi in triacetin (n=3-4). Injection force as a function of particle concentration is fit to a particle jamming model. **d. i)** SEC trace of fresh hIgG control and hIgG formulated in histidine buffer (100 mg/mL), glycine buffer (100 mg/mL), and triacetin (500 mg/mL) after 4 days of stressed aging at 50 °C and **ii)** corresponding high molecular weight fraction for each formulation following stressed aging. **e)** Comparative volume of administration for bolus injection and UHC protein suspension. **f. i)** Representative IVIS images demonstrating subcutaneous absorption of fluorescently-tagged hIgG administered via a PBS bolus injection or a UHC protein suspension. **ii)** Fluorescent signal in the subcutaneous space is fit to a single phase exponential decay mode to identify half-life of subcutaneous absorption (n=5). **iii)** Comparative half-life of subcutaneous absorption for hIgG administered via PBS bolus injection or UHC protein suspensions formulated with triacetin (n=5). **g)** Comparative volume of administration for bolus injection and UHC protein suspension. **h. i)** hIgG serum concentration in human FcRn transgenic mice following hIgG administered via a PBS bolus injection or a UHC protein suspension (n=5-6). Corresponding **ii)** Cmax, **iii)** serum half life, and **iv)** bioavailability. Bar graph in 6d shows mean ± SD, All other plots show mean ± SEM. Statistical comparisons between two groups were made using a two-tailed Student’s t-test with statistical significance as p<0.05.

Using the 70 Triacetin: 30 DMAc non-solvent mixture, the injection force was measured for a series of BSA-MoNi microparticle suspensions as a function of BSA concentration. To ensure accuracy of reported BSA concentrations, the density of the BSA-MoNi microparticles was measured using a pycnometer and found to be 1.31 g/cm^3^. The expected BSA concentration calculated using this density value was corroborated by evaluating the BSA concentration by absorbance (nanodrop) after dissolving a known volume of suspension formulation in aqueous media (**Fig. 4c**). Injection force measurements with BSA-MoNi microparticle suspensions in 70 Triacetin: 30 DMAc demonstrated injection of 600 mg/mL BSA formulations with a clinically relevant injection force of 17 N through a 26G, ½ inch needle at 1 mL/min (**Fig. 4d**). Concentration vs injection force data was fit to a modified version of the Krieger & Dougherty particle jamming model (*52, 53*). Using a spherical shape factor, we found that BSA-MoNi microparticles achieved the theoretical maximum packing density for spheres (volume fraction > 0.74) before jamming in these suspensions (**Fig. 4d, SI Fig. 7, Supplementary Discussion**), indicating that these microparticles are behaving as smooth, hard spheres on account of the enrichment of the high Tg MoNi excipient in the surface layer of the microparticles. By contrast, BSA-Tween microparticles spray dried with an equal molar loading of Tween 80 exhibited similar size and surface morphology (**Fig. 4f**), but suspensions comprising these particles exhibited higher injection forces at equal BSA concentrations, onset of particle jamming at much lower BSA concentrations, and lower maximum concentrations that were injectable under relevant conditions (**Fig. 4d,e**).

**Figure 7.**
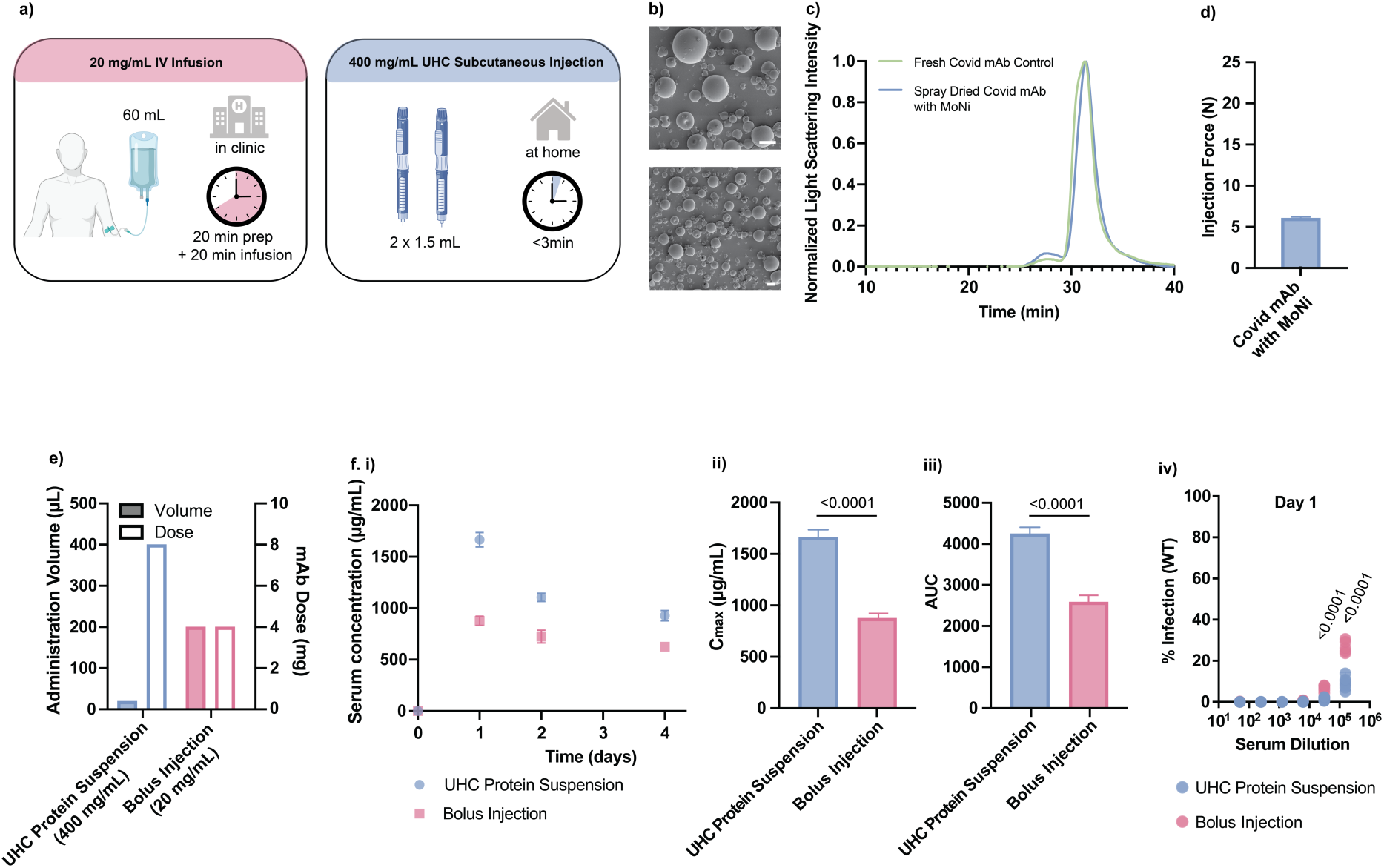
UHC suspension technology can be effectively used to deliver commercial mAb drug product. **a)** Commercial IV antibody drug product (ADP) requires time-intensive delivery in a hospital setting while ADP reformulated as a UHC protein suspension enables delivery with an auto-injector at home. **b)** SEM of spray dried mAb with MoNi particle morphology (scale bars are 10 μm). **c)** SEC trace of fresh mAb control and spray-dried mAb with 5 wt% MoNi. **d)** Injection force of 400 mg/mL mAb microparticles with MoNi in triacetin (n=3). **e)** Comparative volume of administration and mAb dose for bolus injections and UHC protein suspension. **f. i)** mAb serum concentration in human BL6 mice following mAb administered via a bolus injection or a UHC protein suspension (n=5-7). Corresponding **ii)** Cmax, **iii)** bioavailability, and **iv)** day 1 infectivity percent from neutralization. Bar graph in 7d shows mean ± SD, All other plots show mean ± SEM. Statistical comparisons between two groups were made using a two-tailed Student’s t-test with statistical significance as p<0.05.

To demonstrate the translational potential of these UHC protein microparticle suspensions in vivo, we sought to quantify the absorption of protein following SC administration. In these studies, SKH1e mice were injected SC with fluorescently tagged BSA formulated either in a 506 mg/mL triacetin-based suspension comprising MoNi, a 506 mg/mL 70 Triacetin: 30 DMAc suspension comprising MoNi, or a 20 mg/mL PBS solution. Fluorescent images collected from an In Vivo Imaging System (IVIS) were then utilized to study BSA absorption from the site of injection (**Fig. 5a i**). Fluorescent signal over time characterized for the suspension and solution formulations was normalized and fit to a single-phase exponential decay curve to provide half-lives of BSA absorption from the SC space (**Fig. 5a ii**). The half-life of BSA absorption was about 4 hours and statistically insignificant between the three formulations evaluated (**Fig. 5b**), despite a greater than 20-fold difference in required injection volume to achieve similar doses between the UHC suspension and standard solution formulations (**Fig 5c**).

Following these successes with formulation of BSA into UHC suspensions using the MoNi excipient, we applied this technology to a therapeutically relevant protein, human IgG (hIgG). Importantly, while hIgG is the active pharmaceutical ingredient in several dose-limited, commercial drug products (e.g., Cuvitru and Gammagard from Takeda, and Xembify from Grifols) (*2, 54*), hIgG is molecularly comparable with mAb therapeutics in chemical structure, solubility, and size, making it a good model for mAb therapeutics more broadly. Mirroring our previous BSA formulation studies, the hIgG to MoNi ratio in the final spray dried product was fixed at 20:1 (w/w), and the initial feed stock solids concentrations and spray drying parameters were selected to produce particles with desired size and morphology. Spray dried hIgG particles comprising MoNi (5 wt%) were determined to be 5-20 μm in diameter and exhibited a smooth morphology with slightly lower sphericity than the comparable BSA-based particles (**Fig. 6a**). As previously observed with BSA, the inclusion of the MoNi excipient stabilized hIgG during the spray drying process. SEC characterization of fresh hIgG and hIgG spray dried with and without MoNi demonstrated that spray drying hIgG with MoNi resulted in a higher monomer peak fraction and mitigated the formation of high molecular weight aggregates (**Fig. 6b**). Crucially, a control hIgG formulation utilizing 30 wt% trehalose as a stabilizer was found to have a similar stabilizing effect as 5 wt% MoNi, and the addition of 25 wt% trehalose to the 5 wt% MoNi formulation afforded no additional stability benefit to the spray dried hIgG product than MoNi alone (**SI Fig. 8**). In summary, we illustrated that inclusion of only 5 wt% MoNi was sufficient to stabilize hIgG during the spray drying process in a fashion similar to state-of-the-art particle formulations comprising a minimum of 30 wt% trehalose (*55*).

With these high hIgG content microparticles (∼95 wt% protein) prepared with MoNi, we formulated triacetin-based suspensions and measured injection forces as a function of hIgG concentration. In these studies, we used a particle density of 1.35 g/cm^3^, measured by pycnometry, to determine formulation composition. Injection force measurements of 450 mg/mL hIgG formulations of hIgG- MoNi microparticles in triacetin exhibited an injection force of 17 N through a 26G, ½ inch needle (**Fig. 6c**). The concentration vs injection force data for these suspensions were again fit to a modified version of the Krieger & Dougherty particle jamming model (*52*) using a spherical shape factor (**Fig. 6c**). hIgG-MoNi suspensions exhibited jamming at a volume fraction of 0.67 indicating microparticles are behaving as smooth, hard spheres on account of the enrichment of the high- Tg MoNi excipient in the surface layer of the microparticles. Furthermore, similar to our observations with BSA-MoNi microparticle suspensions, the injection force of hIgG-MoNi microparticle suspensions could be further reduced by altering the non-solvent composition (**SI Fig. 9**).

We then assessed the stability of the hIgG protein in suspension through stressed aging assays. In these studies, hIgG-MoNi microparticle suspensions were prepared at 500 mg/mL protein in triacetin. To directly compare to commercial IgG drug products (*2, 54*), aqueous control solutions were prepared with hIgG at 100 mg/mL in histidine or glycine buffers. Formulations were exposed to stressed aging conditions (50 °C and shaking) for one week, and IgG stability at multiple timepoints was assessed by SEC (**Fig. 6d, SI Fig. 10**). hIgG-MoNi suspensions in triacetin showed minimal aggregation at all timepoints and showed lower high molecular weight aggregate formation than aqueous controls. Furthermore, in additional stressed aging studies, we evaluated the stability of the hIgG-MoNi microparticle suspensions in triacetin and aqueous controls through multiple freeze-thaw cycles. The hIgG-MoNi suspension was unchanged following 10 freeze-thaw cycles, while aqueous controls exhibited an increase in high molecular weight species (**SI Fig. 11**). These studies indicated that formulation of the hIgG with MoNi in these microparticle suspensions improved antibody stability.

With UHC hIgG-MoNi suspensions in hand, we then sought to evaluate the SC absorption kinetics of the hIgG from the suspension and a clinically relevant solution formulation. Again, SKH1e mice were injected SC through 27-gauge insulin needles with equal doses of fluorescently tagged hIgG formulation either in a 300 mg/mL triacetin-based suspension formulation comprising MoNi or in a 100 mg/mL PBS solution (**Fig. 6e**). This higher-concentration IgG solution was chosen to be representative of standard aqueous IgG solution drug products. Fluorescent IVIS images were collected over time following administration to study the kinetics of hIgG absorption from the site of injection (**Fig. 6f i**). Fluorescent signal over time for each formulation was normalized and fit to a single-phase exponential decay curve to provide a half-life of hIgG SC absorption (**Fig. 6f ii**). The half-life of hIgG SC absorption was about 9 hours (**Fig. 6f iii**), in agreement with previous literature values (*56*). Similar to our previous observations with BSA-based suspensions, the half-life of absorption was statistically insignificant between the UHC hIgG-MoNi suspension and solution formulations despite a 3-fold difference in the injection volume required for equivalent doses (**Fig. 6e**). To further investigate the in vivo pharmacokinetics of these UHC hIgG-MoNi suspension and solution formulations, hFcRn transgenic mice were injected SC with equal doses of hIgG in either a 350 mg/mL triacetin-based suspension comprising MoNi or a 100 mg/mL PBS solution (**Fig. 6g**). Serum was collected for 9 weeks, and hIgG serum concentrations were quantified by ELISA (**Fig. 6h i**). Again, values for Cmax (**Fig. 6h ii**), serum half-life (**Fig. 6h iii**), and bioavailability (**Fig. 6h iv**) were statistically insignificant between the two formulations despite the greater than 3-fold difference in injection volume (**Fig. 6g**).

Following these successes formulating hIgG into UHC suspensions using the MoNi excipient, we applied this technology to an infectious disease mAb hindered by critical dose limitations. mAbs are useful for pre-exposure or post-exposure prophylaxis for infectious diseases such as SARS-CoV-2. Indeed, Regeneron has commercialized a highly potent anti-spike mAb cocktail for neutralization of SARS-CoV-2 variants (*57*). This mAb cocktail requires ∼1200 mg of mAb to be delivered intravenously in a hospital setting at a relatively low solution formulation concentration of 20 mg/mL (**Fig. 7a**). Here, we demonstrate reformulation of a 20 mg/mL human anti-covid mAb (ID Biologics) as a UHC suspension at 400 mg/mL amenable to subcutaneous administration (**Fig. 7a)**, enabling relevant doses to be delivered by autoinjector in two 1.5 mL subcutaneous injections.

Mirroring our previous hIgG formulation studies, the mAb to MoNi ratio in the final spray dried product was fixed at 20:1 (w/w), and the initial feed stock solids concentrations and spray drying parameters matched those used for hIgG. Spray dried mAb particles comprising MoNi (5 wt%) were determined to be 5-20 μm in diameter and exhibited a smooth spherical morphology (**Fig 7b**). Again, the inclusion of the MoNi excipient stabilized the mAb during the spray drying process. SEC characterization of fresh mAb and mAb spray dried with MoNi demonstrated preservation of the monomer peak and no high molecular weight aggregates (**Fig. 7c**). With these high mAb content microparticles (∼95 wt% protein) prepared with MoNi, we formulated triacetin-based suspensions. We again used a particle density of 1.35 g/cm^3^, measured by pycnometry, to determine suspension formulation composition. Injection force measurements of mAb-MoNi microparticle suspensions in triacetin at 400 mg/mL mAb exhibited an injection force of 6 N through a 26G, ½ inch needle (**Fig. 7d**). This injection force is well below the injection force threshold of 25 N for commercial autoinjectors. Furthermore, these suspensions were stable in triacetin and demonstrated comparable stressed aging profiles to that of hIgG (**SI Fig. 12**).

To investigate the in vivo pharmacokinetics of the UHC mAb-MoNi suspension, BL6 mice were injected SC with either 20 µL of a triacetin-based mAb-MoNi suspension comprising 400 mg/mL protein or 200 µL (n.b., this is the maximum achievable subcutaneous administration volume) of a 20 mg/mL mAb solution (**Fig. 7e**). As such, mice that received UHC mAb suspensions received two times the mAb dose in one-tenth the injection volume. Serum was collected for one week (at which point the onset of anti-drug antibody responses was observed), and anti-SARS-CoV-2 mAb serum concentrations were quantified by ELISA with SARS-CoV-2 receptor-binding domain (RBD) coated plates (**Fig. 7f i**). Mice that received UHC mAb-MoNi suspensions had statistically greater values for Cmax (**Fig. 7f ii**) and bioavailability (**Fig. 7f iii**) than mice receiving bolus injections, and these values were commensurate with dosing. Additionally, comparative neutralization of serum samples at 24 hours after adjusting for dosing demonstrated mAb delivered in UHC mAb suspensions retains neutralization ability (**Fig. 7f iv**). In summary, we demonstrate that human mAb drug products can be successfully reformulated into UHC protein microparticle suspensions, enabling subcutaneous delivery of therapeutic doses that were previously inaccessible.

## 3. Discussion

In this work we leverage a glassy surfactant called MoNi, which is a polyacrylamide derivative with a Tg exceeding that of high Tg sugars such as trehalose (140 °C compared to 106 °C), as an efficient stabilizer for formulating protein therapeutics into readily injectable UHC suspensions. We applied this approach to improve the administration of antibodies, the dominant class of biotherapeutics, which require high concentration formulations to achieve therapeutically relevant doses in a SC injection. Currently, clinically approved products have relied on unstable and burdensome, multi-injection regimes of liquid formulations or the addition of an enzyme (e.g., hyaluronidase) as a secondary active biologic to degrade the SC tissue to enable administration of high-volumes (*11*). Both liquid-based formulation approaches are prone to aggregation of the active biopharmaceutical, limiting shelf-life and global accessibility of these crucial drug products. Recently, formulations of spray-dried protein microparticles suspended in a non-solvent have been developed to enhance protein stability, but current formulation approaches fail to meaningfully increase injectable doses over liquid formulations (typically <300 mg/mL) due to poor microparticle mechanical properties (i.e., low Tg surface properties) and excessively high (>30 wt%) additive concentrations (*34–36*). We hypothesized that a glassy surfactant such as MoNi, with a high Tg (140 °C) and the ability to stabilize protein drugs against harsh accelerated aging conditions, could dramatically improve injectability of these protein suspensions by yielding mechanically robust microparticles at minimal additive concentrations and exceptionally high protein loadings (95 wt%).

It has been previously demonstrated that addition of surface-active excipients, including surfactants like Tween 20 or Tween 80, to aqueous formulations of proteins prior to spray drying helps to maintain drug bioactivity and mitigate protein aggregation through the spray drying process. These surfactants have been shown to be better stabilizers in this context than sugars because they are amphiphilic and preferentially partition to the interfaces formed as the aqueous feed stocks are aerosolized into droplets during spray drying, preventing protein denaturation and aggregation at these air-water interfaces (*37, 39, 40, 58*). These surfactants can also improve spray dried particle morphology. Unfortunately, the amount of these PEG-based surfactants that can be used in these formulations is limited (typically < 0.3 wt%) on account of the low Tg of the excipients (approximately -65 °C), which negatively impacts the surface mechanical properties of the resulting microparticles (*59*). As friction interactions between particles significantly increases the viscosity of suspensions and can lead to paste-like formulations at high particle loadings, the poor mechanical properties induced by the surface enrichment of standard PEG-based surfactants yields suspension formulations that are uninjectable at high protein concentrations. For these reasons, these formulations require inclusion of high Tg (>100 °C) additives, which are typically sugars such as trehalose, at quantities exceeding 30 wt% to yield mechanically robust protein microparticles and suspensions that are both stable and injectable under relevant conditions (*34–36*). Yet, the high loading of these excipient solids within the microparticles severely limits the actual protein content achievable in the resulting suspension formulations.

To address these shortcomings of previous technologies, we replaced these surfactant and sugar additives with a single glassy surfactant additive, MoNi, during the spray drying process. We found that incorporation of MoNi not only stabilized proteins during the spray drying process, but the resulting protein microparticles comprising MoNi could be formulated as readily injectable suspensions at exceptionally high protein contents (exceeding 500 mg/mL for BSA and exceeding 400 mg/mL for human antibodies). We demonstrated that these UHC suspensions could be injected through clinically relevant needles (26G and 27G) with low injection forces relevant to standard pen autoinjector devices (<25 N). Additionally, experiments in mice showed that UHC suspension formulations of a model protein BSA and two antibody-based drug products (hIgG and an anti-SARS-CoV-2 mAb) dramatically reduced required injection volumes without altering pharmacokinetics, protein structure, or efficacy. In summary, we demonstrate that the high Tg excipient MoNi enables stable and readily injectable protein formulations at the highest reported protein concentrations to date (*30, 34, 35*).

The MoNi-containing protein microparticles reported in this work were generated using a standard commercial laboratory-scale spray dryer without fully optimized parameters. We believe that further optimization of the spray drying process parameters and utilizing industrial-scale spray dryers is likely to enable the development of even higher protein concentration suspensions. Given that spray drying is commonly used in the pharmaceutical industry, and there are numerous licensed biopharmaceutical drug products which already utilize particle-based suspension formulations (e.g., Byetta) (*60*), we believe clinical translation of the protein-MoNi microparticle suspension technology we report here is highly feasible.

Additionally, new formulation technologies enabling high-concentration, low-volume antibody delivery with less reliance on the cold-chain will be essential to facilitating global access to critical biopharmaceutical drugs. While mAbs are one of the most effective therapeutic modalities for treating many life-threatening conditions, the majority of approved mAb drug products require high doses (>100 mg per dose) for therapeutic efficacy (*61*). Between 1998 and 2021, the US FDA approved 27 solution-based high-concentration antibody products (>100 mg/mL) that can be administered by SC injection (*54*). Unfortunately, stability and viscosity limitations mean these drug products are typically limited to mAb concentrations of only 200 mg/mL or lower, and many of these approved mAb drug products still require multiple injections to achieve therapeutically relevant doses (*54*). As large SC injection volumes are often associated with pain, the widely accepted maximum volume for SC injection is 1.5 mL (*62, 63*). Indeed, many pen autoinjector devices have been engineered to deliver this volume per administration. As currently formulated, only 8 of the 27 SC injectable high-concentration mAb drug products (30% of these drugs) can be administered with an injection volume of 1.5 mL or less and achieve therapeutic doses (**SI Fig. 13a i**) (*54*). In this work, we demonstrate successful reformulation of commercial low-concentration human antibody drugs into UHC protein suspension at >400 mg/mL. If our protein microparticle suspension technology were applied to reformulation of current mAb drug products, 22 of 27 (>80%) could be delivered as a single 1.5 mL injection (**SI Fig. 13a ii, SI Fig. 13b**).

Overall, the protein formulation platform we describe here both improves the shelf stability and dramatically increases achievable formulation concentrations of clinically relevant biologics. These advancements are highly distinguishing, and as such this technology has the potential to significantly improve accessibility and greatly reduce patient burden associated with these critical therapeutics.

## 4. Materials and Methods

### 4.1 Experimental Design

The purpose of these experiments was to leverage a unique copolymer excipient to enable ultra-high-concentration biologic formulations. The number of experimental repeats/replicates is listed in the figure legends. The initial sample sizes for in vivo experiments was based on Mead’s resource equation. Animals were cage blocked and were only excluded if they were known to be incorrectly dosed at the start of the experiment. Statistical methods used are listed below and also in the figure legends. The investigators were not blinded for these experiments.

### 4.2 Materials

All reagent grade materials and solvents were purchased from Sigma Aldrich or Fisher and used as received. Alexa-647-NHS was purchased from Lumiprobe. Slide-A-Lyzer dialysis cassettes (2 kDa MWCO) from Thermofisher were used for polymer purification. BSA (A2153-50G, CAS-No:9048-46-8) was purchased as a lyophilized powder from Sigma Aldrich. Human IgG (Cat No: 340-21, Lot: 07J4627) was purchased as a lyophilized powder from Medix Biochemica. HyPure™ Cell Culture Grade Water was purchased from Cytiva. Phosphate Buffered Saline (10010–023) was purchased from Gibco. Syringes used for injection force measurements are Fisherbrand 1mL plastic leur lock syringes (Cat. No: 14955464). Syringes used for formulating protein suspensions with non-solvent were Thermo Scientific 5mL leur slip plastic syringe (Cat. No: S7510-5). Needles used for injection force measurements were BD PrecisionGlide™ needles (26G, ½ in, Ref: 305111). In vivo protein suspension delivery was performed using BD Insulin Syringes with BD Micro-Fine™ IV Needles (27G, 12.7mm).

### 4.3 Formulating UHC protein suspensions

Protein suspensions were formulated by combining spray dried protein microparticles with a non-solvent.

#### 4.3.1 MoNi synthesis and characterization

Polymers were synthesized and characterized according to previously published methods.(*20*) In brief, to a 20 mL scintillation vial 2-cyano-2-propyl dodecyl trithiocarbonate (CDPT, 259 mg, 0.75 mmol), 2,2′-azobis(2-methyl-propionitrile) (AIBN, 24.6 mg, 0.15 mmol), 4-acryloylmorpholine (Mo, 3.85 g, 27.3 mmol), N-isopropylacrylamide (Ni, 1.15 g, 10.2 mmol), and dimethyl formamide (DMF, 9.0 mL). The vial was capped with a PTFE septa and sparged for 20 minutes. The polymerization was conducted at 65 °C for 18 h to a conversion above 95% as determined by H^1^ NMR (Bruker Neo-500 MHz). Polymers were purified by precipitating 3 times in 75:25 ether:hexane mixture. Then the end group was removed by addition of the polymer to a 20-fold excess AIBN (4.1 g, 25 mmol), lauroyl peroxide (LPO, 996 mg, 2.5 mmol) and DMF (45 mL) in a 200 mL round bottom flask and the reaction was heated to 90 °C for 24 h. Afterwards, excess AIBN was removed by precipitation and the polymer was dialyzed against DI water for 48 h to prepare the final product.

To characterize the final MoNi polymer we determined the molecular weight (Mn SEC) and dispersity using an RI detector and polymethylmethacrylate standards. The running solvent was N,N-dimethylformamide (DMF) with 1 g/L LiBr (flow rate: 1 mL/min) heated to 50 °C and samples were prepared at 5 mg/mL. Separation was done through two Jordi Labs Resolve Mixed Bed Low Divinylbenzene (DVB) columns in series and data was collected by a Dionex Ultimate 3000 Variable Wavelength detector and RefractoMax521 RI detector. The RI traces were normalized and areas under the curves for the 310 nm absorbance signals were calculated with Prism 10.

#### 4.3.2 Spray drying BSA microparticles

BSA feed solutions were prepared by dissolving lyophilized BSA in cell grade water at 2 wt% (20 mg/mL). BSA solutions dissolved at room temperature for one hour prior to sterile filtering using a 0.2 μm sterile filter. After sterile filtering, 7kDa MoNi was added to the feed solution at a concentration of 0.1wt% (1 mg/mL). Feed solutions were stored on ice prior to spray drying.

Samples were spray dried using a Buchi B-290 Mini Spray Dryer equipped with a high-performance cyclone. Samples were spray dried using an inlet temperature of 150C (outlet ∼67C), aspirator pressure of 40 mm, and a pump rate of 20% (6 mL/min). Collected powder was transferred to a 50mL falcon tube and stored at 4 °C with desiccant.

#### 4.3.3 Spray drying BSA microparticles with Tween 80

BSA feed solutions were prepared by dissolving lyophilized BSA in cell grade water at 2 wt% (20 mg/mL). BSA solutions dissolved at room temperature for one hour prior to sterile filtering using a 0.2 μm sterile filter. After sterile filtering, Tween 80 was added to the feed solution at a concentration of 0.0176 wt% (0.176 mg/mL). Tween 80 concentration was chosen to add equal moles of MoNi and Tween 80 to the spray drying feed solution. Feed solutions were stored on ice prior to spray drying.

Samples were spray dried using an inlet temperature of 150 °C (outlet ∼67C), aspirator pressure of 40 mm, and a pump rate of 20% (6 mL/min). Collected powder was transferred to a 50mL falcon tube and stored at 4 °C with desiccant.

#### 4.3.4 Spray drying hIgG microparticles

hIgG feed solutions were prepared by dissolving lyophilized hIgG in cell grade water at 10 wt% (100 mg/mL). hIgG solutions dissolved at 4 °C for four hours prior to sterile filtering using a 0.2 μm sterile filter. The sterile-filtered product was then dialyzed against cell grade water overnight using a 30K MWCO Slide A Lyzer Dialysis Cassette to remove residual salts in the lyophilized product. After dialysis, the hIgG concentration of the solution was quantified by nanodrop, and the volume of the solution was measured using a graduated cylinder. The hIgG solution was then diluted 50 mg/mL hIgG using cell grade water. 7kDa MoNi was added to the feed solution at a concentration of 0.25 wt% (2.5 mg/mL). Feed solutions were stored on ice prior to spray drying.

Samples were spray dried using an inlet temperature of 80 °C (outlet ∼53C), aspirator pressure of 40 mm, and a pump rate of 1 mL/min. Collected powder was transferred to a 50mL falcon tube and stored at 4 °C with desiccant.

#### 4.3.5 Spray drying covid mAb microparticles

Human anti-SARS-CoV-2 mAb was purchased from ID Biologics. The mAb drug product is formulated at 20 mg/mL in a non-disclosed buffer. The mAb drug product was first buffer exchanged into cell grade water by centrifugal filtration using an Amicon® Ultra-15 30,000 MWCO device spun at 4000 x g. Buffer exchange was performed by 5 repeated cycles of 10-fold concentration and dilution with cell grade water. Following buffer exchange, the concentrated mAb was diluted to 100 mg/mL and sterile filtered using a 0.2 μm sterile filter. After sterile filtering, the concentration of mAb in the solution was quantified by nanodrop, and the volume of the solution was measured using a graduated cylinder. The mAb solution was then diluted 50 mg/mL mAb using cell grade water. 7kDa MoNi was added to the feed solution at a concentration of 0.25 wt% (2.5 mg/mL). Feed solutions were stored on ice prior to spray drying.

Samples were spray dried using an inlet temperature of 80 °C (outlet ∼53C), aspirator pressure of 40 mm, and a pump rate of 1 mL/min. Collected powder was transferred to a 50mL falcon tube and stored at 4 °C with desiccant.

#### 4.3.6 Microparticle imaging

Microparticle morphology was characterized by scanning electron microscopy (SEM). Samples were grounded to an aluminum pin stub using double-sided conductive copper tape. A 5.0 nm thick layer of pure gold was deposited onto the samples using a Leica ACE600 Vacuum system. SEM analysis was performed using the FEI Magellan 400 XHR Scanning Electron Microscope at 5.00 kV and high vacuum in field-free mode.

#### 4.3.7 Microparticle density measurements

Microparticle density was measured using an AccuPyc 1330. A known sample mass of spray dried powder between 200 and 300 mg was measured into a small sample cell with a volume of 1 cm^3^. Powder volume was measured over 999 cycles. The known mass, as measured by analytical balance, and the average sample volume was used to calculate powder density.

#### 4.3.8 Formulating suspensions

Suspensions were formulated by combining a known mass of spray dried microparticles with a known volume of non-solvent. Protein concentration in mg/mL was determined by assuming total volume encompassed non-solvent volume as well as spray dried particle volume. When calculating suspensions concentration, the spray dried particle density was taken from pycnometry experiments.

To minimize non-solvent evaporation when preparing suspensions, spray dried microparticles were added to the barrel of a 6mL luer slip syringe. The mass of spray dried particles was measured using an analytical balance. The desired volume of non-solvent or non-solvent combination was added to the syringe barrel through the syringe tip using a p200 pipette. After the syringe was capped, and the protein suspension was mixed inside the syringe using a vortex for 5 minutes or until all powder was fully dispersed. Protein suspensions were transferred from 6mL leur slip syringes to the alternative desired syringe (1 mL leur lock syringes or insulin syringes) by back loading for force of injection experiments or animal experiments respectively.

### 4.4 Characterizing flow properties of UHC protein suspension

The flow properties of protein suspensions were characterized through rheology and injection force measurements.

#### 4.4.1 Rheological characterization

Rheological testing was performed using a stress-controlled TA Instruments DHR-2 rheometer. Rheology of solid-like formulations (BSA without MoNi) was performed at 25 °C using a 20 mm diameter serrated parallel plate at a 500 µm gap. Rheology of liquid-like formulations (BSA with MoNi) was performed at 25 °C using a 40 mm cone geometry with a 50 µm gap. Frequency sweeps were performed at a strain of 1% within the linear viscoelastic regime. Flow sweeps were performed from high to low shear rates with steady state sensing.

#### 4.4.2 Injection force measurements

Force of injection was quantified by measuring the force required to inject a protein suspension through a known needle gauge at a known flow rate using a syringe of known barrel dimensions. A force sensor was built that encompassed a load cell (FUTEK LLB300 50 lb Subminiture Load Button (Model #: LLB300, Item #: FSH03954, Serial #: 705242)) attached to a syringe pump (KD Scientific Syringe Pump (Model #: LEGATO 100, Catalog #: 788100, Serial #: D103954)). An Omega Engineering Platinum Series Meter (Model #: DP8PT, Serial #: 18110196) was used to translate load cell resistance measurements to force values in Kg. The load cell was calibrated prior to measuring injection force (**SI Fig. 14**). A lab view program records the forces measured throughout the duration of an injection experiment and displays a graph of injection force over time.

Injection force experiments were performed as follows. A 1mL Thermo Fisher luer lock syringe with the desired needle gauge was loaded into the syringe pump. The syringe pump height was adjusted so that the load button of the force sensor was in contact with the end of the syringe plunger. The initial force was at or very close to 0 Kg. The appropriate syringe barrel dimensions as well as desired flow rate and injection volume were then selected. The syringe pump moved at the programmed rate injecting protein suspension through the attached needle. The force sensor coupled with the Omega unit measured the force required to inject the protein suspension at the desired flow rate. A lab view program recorded the forces measured throughout the duration of an injection experiment and displayed a graph of injection force over time. Force of injection was quantified by subtracting the average initial force (background) from the average plateau injection force. Injection force in Kg was converted to injection force in Newtons by multiplying by 9.81.

### 4.5 Characterizing protein stability

Protein stability before and after spray drying was characterized by SEC. Protein suspension stability was characterized by SEC as well as by comparative injection force.

#### 4.5.1 SEC Characterization

The SEC trace of protein samples was determined with the ASTRA software package (Wyatt Technology Corporation) after passing a 5 mg/mL protein sample through a size-exclusion chromatography column (Superose 6 Increase 10/300 GL) in a mobile phase of PBS with sodium azide at 25 °C and a flow rate of 0.5 mL/min. Detection consisted of a Optilab T- rEX (Wyatt Technology Corporation) refractive index detector operating at 658 nm and a TREOS II light scattering detector (Wyatt Technology Corporation) operating at 659 nm. A d*n*/d*c* value of 0.185 was used for BSA and IgG samples. To directly compare the stability of spray dried protein samples, SEC traces were normalized to the height of the monomer peak.

#### 4.5.2 Methodology for BSA stressed aging

BSA protein suspensions were prepared at 520 mg/mL in triacetin using spray dried BSA particles with and without 5wt% MoNi. A BSA aqueous control was prepared by dissolving fresh, lyophilized BSA at 20 mg/mL in PBS. All samples were stored in parafilmed 8mL scintillation vials. A 500 mL beaker of water was heated to 60 °C using a temperature controlled hot plate. The sample files were submerged in the 60 °C water bath so that the protein sample volume was fully beneath the water line. Samples were heated at 60 °C for 30 minutes to encourage protein degradation. Following stressed aging, samples were redissolved in PBS at 5 mg/mL, and protein stability was assessed through SEC.

#### 4.5.3 Methodology for hIgG and human mAb stressed aging

hIgG microparticle suspensions were prepared at 500 mg/mL protein in triacetin. Aqueous control solutions were prepared with hIgG at 100 mg/mL in 0.25M histidine or 0.25M glycine buffers. Formulations were exposed to stressed-aging conditions (50 °C incubator and shaking at 160 RPM) for one week, and IgG stability at multiple timepoints was assessed by SEC. Human mAb microparticle suspensions were prepared at 400 mg/mL protein in triacetin, and stability through stressed aging was accessed in the same manner.

#### 4.5.4 Methodology for hIgG stability through freeze-thaw cycles

hIgG microparticle suspensions were prepared at 500 mg/mL protein in triacetin. Aqueous control solutions were prepared with hIgG at 100 mg/mL in 0.25M histidine or 0.25M glycine buffers. Formulations were freeze-thawed up to 10 times, where each freeze thaw cycle consisted of allowing the sample to reach -80 °C and subsequently allowing the sample to warm to room temperature (20 °C). IgG stability at multiple freeze-thaw cycle timepoints was assessed by SEC.

#### 4.5.5 UHC protein suspension storage and injection force measurements

BSA protein suspensions with 5wt% MoNi were prepared at 520 mg/mL in triacetin. Protein suspensions were loaded into a 1 mL Thermo Fisher luer slip syringe and capped with a BD 26G ½ inch needle wrapped with parafilm to limit solvent evaporation. Injection force for the 520 mg/mL protein suspension was measured on day 0 and again on day 35. The syringe was stored horizontally at 23 C.

### 4.6 In vivo delivery of UHC protein suspensions

Animal studies were performed with the approval of the Stanford Administrative Panel on Laboratory Animal Care (APLAC-32109) and were in accordance with National Institutes of Health guidelines.

#### 4.6.1 IVIS Imaging of BSA delivery

Fluorescently tagged BSA microparticles were obtained by spray drying AF647-BSA with untagged BSA at a ratio of 1:500. MoNi was kept at 5 wt% in the final particle formulation. Fluorescently tagged BSA microparticles were then further diluted with untagged BSA microparticles with MoNi at a ratio of 1:5 resulting in a final ratio of 1:2500 AF647-BSA: untagged BSA. Protein suspensions in triacetin and 70 triacetin: 30 DMAc were formulated as described above. A bolus control consisted of BSA at a ratio of 1:2500 AF647-BSA: untagged BSA dissolved in PBS at 20 mg/mL.

Female SKH1-Elite Mice were each given subcutaneous injections of either 8.8 μl of 506 mg/mL fluorescently-tagged BSA protein suspension (in triacetin or 70 triacetin: 30 DMAc) or 200 uL of 20 mg/mL fluorescently tagged BSA in PBS. Protein suspension injections were administered with a 50 uL hamilton syringe with a 26G ½ inch needle. Bolus injections were administered with a 1 mL luer lock syringe with a 26G ½ inch needle. The subcutaneous injection sites of animals were imaged using the IVIS (Lago) over a series of time points spanning two days. When imaging, mice were anesthetized with isoflurane gas and imaged with an exposure time of 2 s, excitation wavelength of 600 nm, and emission wavelength of 670 nm (binning, medium; F/stop, 1.2). Total radiant efficiency [(photons/s)/(μW/cm2)] was quantified using an equal-sized region of interest surrounding the injection site. Fluorescence intensity at each time point was normalized to the maximum fluorescent intensity and normalized fluorescence intensity values for each mouse (n = 3-5) were fit to a single exponential decay model, and half-lives were acquired and averaged using GraphPad Prism.

#### 4.6.2 IVIS imaging of hIgG delivery

hIgG was fluorescently labeled with Alexa-647-NHS from lumiprobe at 5% wt ratio. In brief, 50 mg dry hIgG was dissolved in 10 mL of PBS and 500 uL of a 5 mg/mL DMSO stock solution of Alexa-647-NHS was added to the solution. The reaction proceeded for 24 h at room temperature and the free dye was removed with 10 kDa MWCO amicon spin filter.

Fluorescently tagged hIgG microparticles were obtained by spray drying AF647-tagged hIgG with untagged hIgG at a ratio of 1:20. MoNi was kept at 5 wt% in the final particle formulation. Fluorescently tagged hIgG microparticles were then further diluted with untagged hIgG microparticles with MoNi at a ratio of 1:100 resulting in a final ratio of 1:2000 AF647-tagged hIgG: untagged hIgG. Protein suspensions in triacetin were formulated as described above. A bolus control consisted of hIgG at a ratio of 1:2000 AF647-tagged hIgG: untagged hIgG dissolved in PBS at 100 mg/mL.

Female SKH1-Elite Mice were each given subcutaneous injections of either 20 μl of 300 mg/mL fluorescently-tagged hIgG protein suspension in triacetin or 60 uL of 100 mg/mL fluorescently tagged hIgG in PBS (n=5). Protein suspension and bolus injections were both administered subcutaneously into the flank of the mouse with an insulin syringe with a 27 gauge needle. The subcutaneous injection sites of animals were imaged using the IVIS (Lago) over a series of time points spanning two days.

IVIS imaging and half-life data analysis were performed as described above. Animals were imaged with an exposure time of 1 s.

#### 4.6.3 In vivo Pharmacokinetics of hIgG delivery

Female scid FcRn-/- hFcRn Tg mice were each given subcutaneous injections of either 30 uL of 350 mg/mL hIgG protein suspension in triacetin or 105 uL of 100 mg/mL fresh hIgG in PBS to result in a uniform protein dose of 10.5 mg of hIgG per mouse. When formulating protein suspensions in triacetin, it was assumed that spray dried powder had a density of 1.35 g/cm^3^ and was 95% hIgG by mass. Protein suspension and bolus injections were both administered subcutaneously into the flank of the mouse. At selected timepoints mice were anesthetized with isoflurane gas, and blood samples were drawn via tail vein. Timepoints included 24 hr, 48hr, 4d, 6d, 11d, 16d, 21d, 28d, 35d, 49d, and 63d. The concentration of hIgG in serum was quantified using an IgG (Total) Human ELISA Kit (ThermoFisher Catalog #BMS2091). Pharmacokinetic readouts included Cmax, serum half-life, and bioavailability. Cmax for each animal was the highest hIgG titer quantified by IgG Total Human ELISA. Serum half-life was quantified by fitting hIgG titers (48h-day 63) for each treatment group to a single-phase exponential decay model, and half-lives were acquired using GraphPad Prism. Bioavailability was quantified by calculating area under the curve (AUC) (0-day 63) for each animal in the PBS bolus and triacetin suspension using GraphPad Prism. Bioavailability of hIgG in each animal was quantified by dividing the AUC of each animal by the average AUC of the PBS bolus group.

#### 4.6.4 In vivo Pharmacokinetics of human mAb delivery

Female BL6 mice were each given subcutaneous injections of either 20 µL of 400 mg/mL covid mAb (ID Biologics) protein suspension in triacetin or 200 µL of 20 mg/mL covid mAb (ID Biologics) drug product. When formulating protein suspensions in triacetin, it was assumed that spray dried powder had a density of 1.35 g/cm^3^ and was 95% mAb by mass. Protein suspension and bolus injections were both administered subcutaneously into the flank of the mouse. At selected timepoints mice were anesthetized with isoflurane gas, and blood samples were drawn via tail vein. Timepoints included 24 hr, 48hr, 4d, 7d, 10d, and 14d. The concentration of mAb in serum was quantified by ELISA with SARS-CoV-2 (2019-nCoV) Spike RBD-His (A435S) Recombinant Protein (Sino Biological) coated plates (at 2 µg/mL). Pharmacokinetic readouts included Cmax and bioavailability. Cmax for each animal was the highest mAb titer quantified by ELISA. Relative bioavailability was quantified by calculating area under the curve (AUC) (0- day 4) for each animal using GraphPad Prism.

#### 4.6.5 Pseudovirion neutralization experiments

Neutralization experiments were performed as described in Xu et al. (*64*). Briefly, SARS-CoV-2 spike-pseudotyped lentiviruses encoding a luciferase-ZsGreen reporter were produced in HEK293F cells by co-transfection of five plasmids. The five plasmids include a packaging vector (pHAGE-Luc2-IRES-ZsGreen), a plasmid encoding the SARS-CoV-2 spike (HDM-SARS2-spike-WT) and three helper plasmids (pHDM-Hgpm2, pHDM-Tat1b and pRC-CMV_Rev1b). 50 mL of cells were diluted to a density of approximately 3-4 × 10^6^ cells/mL. Transfection mixture was prepared by adding five plasmids (50 µg of packaging vector, 17 µg of SARS-CoV-2-encoding plasmid and 11 µg of each helper plasmid) to 5 ml of Expi-free medium, followed by the dropwise addition of BioT transfection reagent (150 µl, Bioland Scientific) with vigorous mixing. After 10 min incubation at room temperature, the transfection mixture was transferred to HEK293F cells. D-glucose (4 g/L, Sigma-Aldrich) and valproic acid (3 mM, Acros Organics) were then added to the cells immediately post-transfection to increase recombinant protein production. The cells were harvested 3-5 days after transfection by spinning the cultures at 300 x g for 5 minutes. The supernatant was filtered through a 0.45-µm filter and 0.5 mL of 1mM HEPES was added to neutralize the pH. Viral stocks were aliquoted and flash-frozen in liquid nitrogen. They were stored at −80 °C and titrated before further use.

Antisera were heat inactivated (56 °C, 30-60 min) before neutralization assays. Neutralization against SARS-CoV-2 WT was analyzed in HeLA-ACE2/TMPRSS2 cells. One day before infection (day 0), cells were seeded at 8,000 cells per well in white-walled, white-bottom, 96-well plates (Thermo Fisher or Greiner Bio-One). On day 1, antisera were serially diluted in D10 media and mixed 1:1 with pseudoviruses for 1-2 hrs at 37 °C before being transferred to cells. The pseudovirus mixture contained SARS-CoV-2 WT, D10 media and polybrene (1:500). Assays were read out with luciferase substrates 2 days after infection by removing the media from the wells and adding 80 uL of a 1:1 dilution of BriteLite in DPBS (BriteLite Plus, Perkin Elmer). Luminescence values were measured using a microplate reader (BioTek Synergy™ HT or Tecan M200). Percent infection was normalized on each plate. Neutralization assays were performed in technical duplicates.

### 4.7 Statistical analysis

Injection force data is reported as means with standard deviation. For in vivo experiments, animals were cage blocked, and Mead’s Resource Equation was used to identify a sample size above that additional subjects will have little impact on power. Normalized fluorescence intensity and half-life of subcutaneous absorption from in vivo experiments are reported as means with standard error. Comparison between two groups was conducted with a two-tailed Student’s t-test. Comparison between greater than two groups was conducted with the Tukey HSD test in JMP. Results were accepted as significant if *p* < 0.05.

## 5. List of Supplementary Materials

Supplementary Figures 1-14 Supplementary Discussion

## Acknowledgements

We appreciate the support of Dr. Jayakumar Rajadas and the Biomaterials and Advanced Drug Delivery (BioADD) Laboratory for allowing us to use their Buchi B-290 spray dryer. We appreciate the support of Alexandra Grande and the Stanford Dorre School of Sustainability Sample Preparation Laboratory for allowing us to use their ball mill. We are grateful to Cynthia M. Ross for assisting with pycnometer experiments. We thank Benjamin Arbaugh for helpful conversations regarding optimizing spray drying parameters. A.N.P. is supported by a Stanford Maternal and Child Health Research Institute postdoctoral fellowship. C.K.J. is supported by a National Science Foundation Graduate Research Fellowship. N.E. is supported by a National Science Foundation Graduate Research Fellowship (#DGE- 2146755). Ashley Utz is supported by Stanford University Medical Scientist Training Program grant T32-GM007365 and T32GM145402.

## 6.1. Author Contributions

C.K.J., A.N.P., and E.A.A. conceived of the idea. C.K.J. and A.P. performed experiments and data analysis. N.E. helped with data analysis. C. D. helped with SEM imaging. A.U. performed neutralization experiments.

