## Supplemental Information for "Glassy Surfactants Enable Ultra-High Concentration Biologic Therapeutics"

**The PDF includes:**
**Supplementary Figures**

### Supplementary Figures

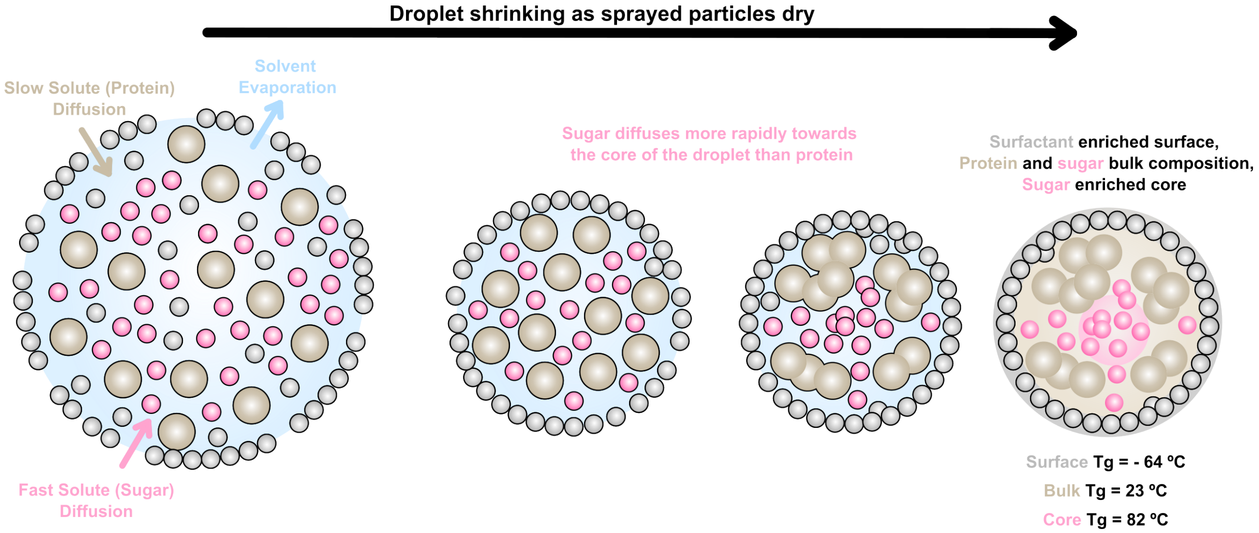

**SI Figure 1. Localization of surfactant and sugar in spray-dried protein particles.** High glass transition temperature (Tg) additives, such as trehalose, are traditionally needed to stabilize the mAbs during the spray drying process and to achieve sufficiently glassy microparticles to form robust injectable suspensions. Tgs were calculated with the Fox equation, based on values of Surfactant (Tween 20, - 64 °C), Sugar (Trehalose, 106 °C) and Protein (-29 °C). Compositions of the core, bulk and surface were approximated from Peclet numbers, diffusion coefficients, and surface activity resulting in 100 wt% surfactant on the surface, 50 wt% protein and 50 wt% sugar in the bulk, and 10 wt% protein and 90 wt% sugar in the core.

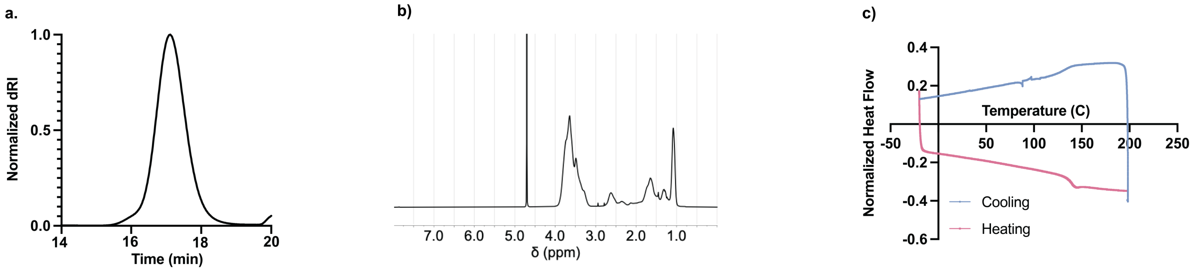

**SI Figure 2. Characterization of MoNi. a)** SEC trace of MoNi in DMF with LiBr. **b)** $^{1}H$ NMR of MoNi in $D_2O$. **c)** DSC of MoNi at a temperature ramp and cooling rate of 10 C/min showing a glass transition temperature between 130 and 140 ºC.

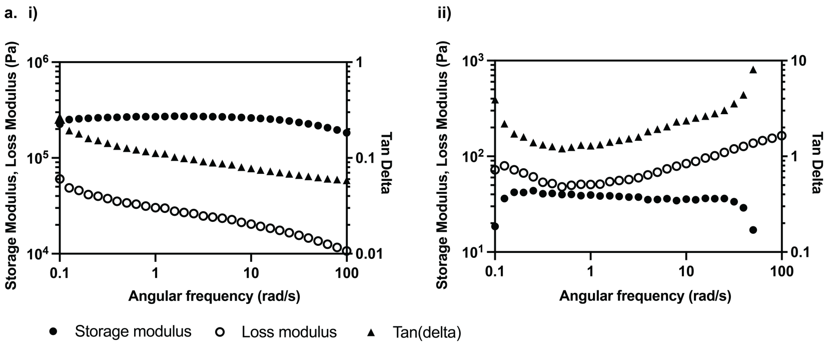

**SI Figure 3. Rheological characterization of spray dried BSA microparticle suspensions. a)** Angular frequency sweep of **i)** 520 mg/mL BSA microparticles resuspended in triacetin and **ii)** 520 mg/mL BSA microparticles containing 5 wt% MoNi resuspended in triacetin.

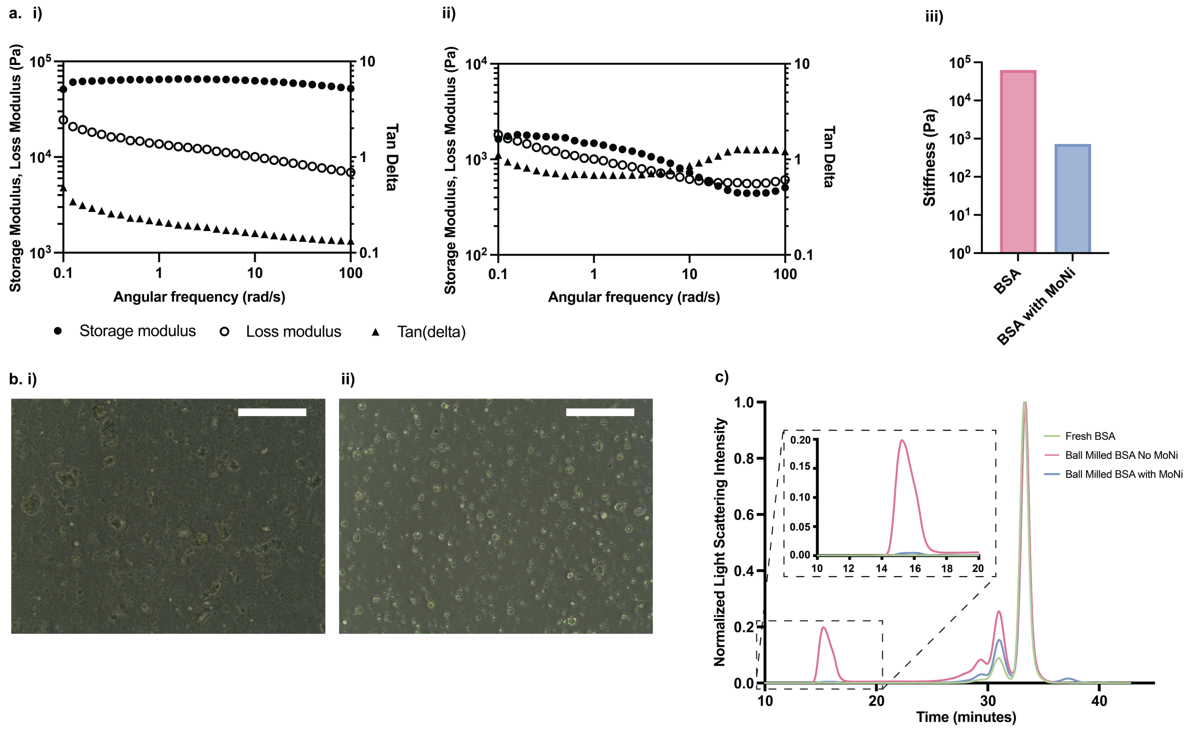

**SI Figure 4. Rheological and stability characterization of ball milled BSA microparticle suspensions.** Ball milling serves as an alternative means to produce protein microparticles. Protein microparticles formed with ball milling on average are larger and more polydisperse than particles formed by spray drying. Additionally, ball milling leads to increased protein aggregation because of high temperatures and mechanical forces. Shown above are an **a)** angular frequency sweep of **i)** 400 mg/mL BSA microparticles resuspended in triacetin, **ii)** 400 mg/mL BSA microparticles containing 5 wt% MoNi resuspended in triacetin, and **iii)** a comparison of the storage modulus of suspensions at 10 rad/sec demonstrating that ball milled BSA with MoNi shows reduced stiffness compared to BSA ball milled without MoNi when formulated in triacetin at equal protein concentration (ball milled particle density assumed to be 1 g/cm$^3$)**. b)** Microscope images of BSA with 5 wt% MoNi microparticles formed by **i)** 15 minutes of ball milling or **ii)** spray drying. Particles are resuspended in sesame oil for improved imaging. **c)** SEC trace of fresh BSA control, 15 minute ball milled BSA without MoNi, and 15 minute ball milled BSA with 5 wt% MoNi. PBS with sodium azide is used as an eluent. Similarly to spray drying, MoNi stabilizes BSA through the ball milling process and results in a final product with substantially reduced high molecular weight aggregates compared to BSA samples ball milled without MoNi.

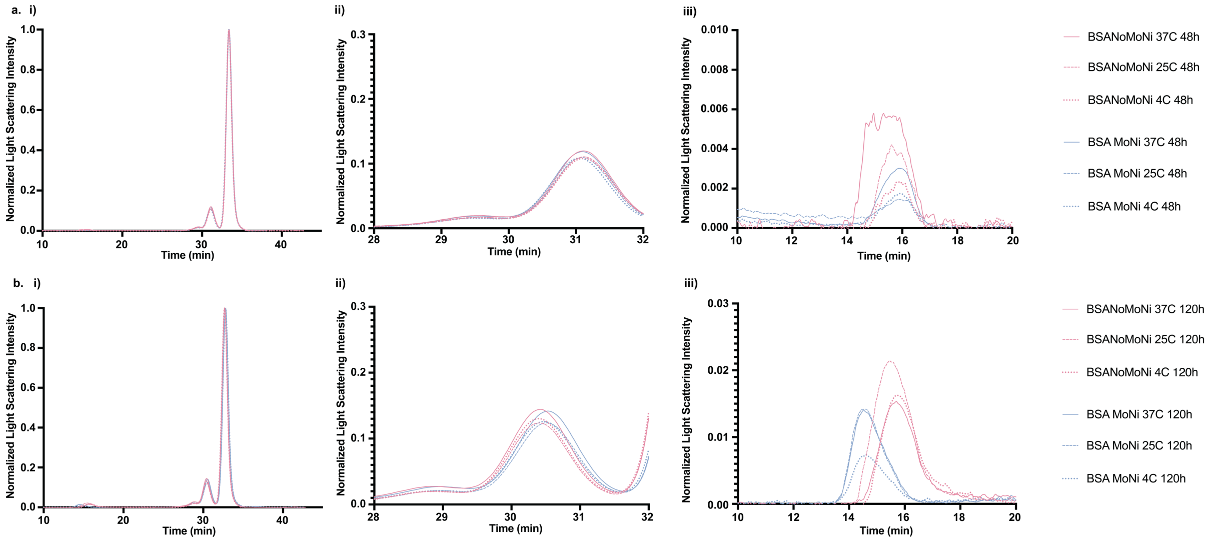

**SI Figure 5. Storage stability of suspensions.** 520 mg/mL BSA formulations in triacetin with and without 5 wt% MoNi were stored for 48 hours and 120 hours at 4C, 25C, and 37 °C to assess stability. **a. i)** Full SEC trace, **ii)** dimer peak, and **iii)** high molecular weight peak of all formulations following 48 hours of storage. **b. i)** Full SEC trace, **ii)** dimer peak, and **iii)** high molecular weight peak of all formulations following 120 hours of storage.

a)

Degree of adverse skin reaction was quantified using the following grading scale:

| Score | Grade | Description |
|---|---|---|
| 0 | None | No swelling, normal pink |
| 1 | Minimal | Slight swelling with indistinct border and/or light pink coloration |
| 2 | Mild | Defined swelling with distinct boarder or bright pink coloration |
| 3 | Moderate | Defined swelling with distinct boarder and bright pink coloration |
| 4 | Major | Small ulceration (<1 mm diameter) along with pronounced swelling (>1mm) or redness |
| 5 | Severe | Large ulceration (>1 mm diameter) along with pronounced swelling (>1mm) or redness |

b)

| | Time | | | | | |
|---|---|---|---|---|---|---|
| Solvent (10uL) | 0 hr | 30 min | 1 hr | 3 hr | 6 hr | 24 hr |
| 100 Triacetin | 0 | 0 | 0 | 0 | 0 | 0 |
| 100 DMAc | 2 | 4 | 2 | 1 | 1 | 1 |
| 50:50 Triacetin: DMAc | 1 | 1 | 0 | 0 | 0 | 0 |
| 70:30 Triacetin: DMAc | 0 | 1 | 0 | 0 | 0 | 0 |
| 75:25 Triacetin: DMAc | 0 | 1 | 0 | 0 | 0 | 0 |
| 80:20 Triacetin: DMAc | 0 | 1 | 0 | 0 | 0 | 0 |
| 100 BA | 2 | 4 | 2 | 1 | 1 | 1 |
| 50:50 Triacetin: BA | 1 | 1 | 0 | 0 | 0 | 0 |
| 70:30 Triacetin: BA | 0 | 1 | 0 | 0 | 0 | 0 |
| 80:20 Triacetin: BA | 0 | 1 | 0 | 0 | 0 | 0 |
| 100 NMP | 2 | 5 | 3 | 2 | 1 | 1 |
| 50:50 Triacetin: NMP | 1 | 4 | 1 | 1 | 1 | 0 |
| 70:30 Triacetin: NMP | 1 | 2 | 2 | 1 | 1 | 1 |
| 80:20 Triacetin: NMP | 1 | 1 | 0 | 0 | 0 | 0 |

**SI Figure 6. Solvent tolerability screen. a)** Grading scale for quantifying degree of adverse skin reaction. **b)** SKH1e mouse skin reaction following subcutaneous administration of 30 µL of solvent blend. These results indicate that triacetin and blends of triacetin with DMAc are well tolerated during subcutaneous injection. Mice administered triacetin alone showed no skin swelling or redness. Although DMAc or BA alone resulted in skin irritation, blends with less than or equal to 50:50 v/v DMAc or BA in triacetin were also well tolerated. These blends demonstrated adverse skin reaction grades of less than or equal to 1 in the first 24 hours after subcutaneous injection. NMP subcutaneous injection resulted in greater skin irritation than DMAc or BA. Tolerability studies demonstrate triacetin, DMAc, and BA are promising non-solvents for *in-vivo* use. Initial in-vivo work focused on triacetin and DMAc due to this blend's improved protein suspension stability and lower injection forces.

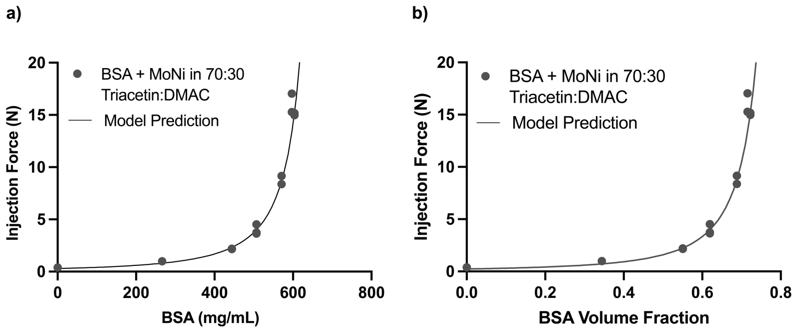

**c)** **Krieger & Dougherty model:**

$$\eta = \eta 0\left(1-\frac{\Phi}{\Phi\text{max}}\right) - [\eta] * \Phi\text{max}$$

**Modified Krieger & Dougherty model:**

$$F_{inj} = F_{inj}0\left(1-\frac{\Phi}{\Phi\text{max}}\right) - [\eta] * \Phi\text{max}$$

| Non-solvent injection force ($F_{inj}0$) | Intrinsic viscosity ([η]) | Max solid fraction (Φmax) |
|---|---|---|
| 0.24 | 2.5 | 0.84 |
| Fixed | Fixed | Fit |

**SI Figure 7. Modeling injection forces of microparticle suspensions. a)** Injection force as a function of concentration for BSA microparticles through a 26G ½ inch needle at 1mL/min. **b)** Injection force as a function of particle volume fraction. **c)** Krieger & Dougherty model (1959) modified for injection force and fit for $\Phi_{max}$.

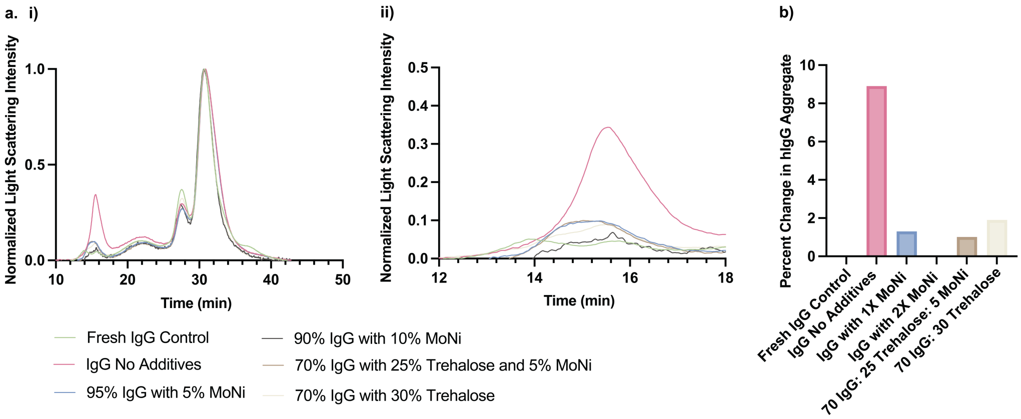

**SI Figure 8. hIgG stability during spray drying. a. i)** Full SEC trace of fresh hIgG control, spray dried hIgG with no additives, spray dried hIgG with 1X MoNi (5 wt%), spray dried hIgG with 2X MoNi (10 wt%), spray dried higG with 25 wt% trehalose and 5 wt% MoNi, and spray dried IgG with 30 wt% Trehalose. **ii)** SEC trace of only high molecular weight aggregate. PBS with sodium azide is used as an eluent. **b)** Percent change in hIgG aggregates due to spray drying, as a percent of total protein content. The baseline for determining percent change was determined by subtracting the starting aggregate percent of the fresh IgG control (3.9%) from all samples.

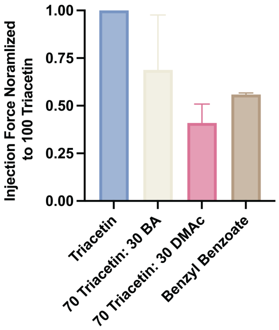

**SI Figure 9. Normalized injection force with alternative non-solvent additives.** Injection of 388 mg/mL and 450 mg/mL hIgG, 5 wt% MoNi suspensions at 1mL/min through 26G ½ inch needles with DMAc, BA and BB non-solvent additives (n=2-5). Like our observations with BSA-MoNi microparticle suspensions, the injection force of hIgG-MoNi microparticle suspensions could be further reduced by altering the non-solvent composition. Using solvent mixtures 70 Triacetin: 30 BA, 70 Triacetin: 30 DMAc, and pure benzyl benzoate, the injection force of hIgG-MoNi microparticle suspensions were reduced by 1.5-fold, 2.4-fold, and 1.8-fold respectively.

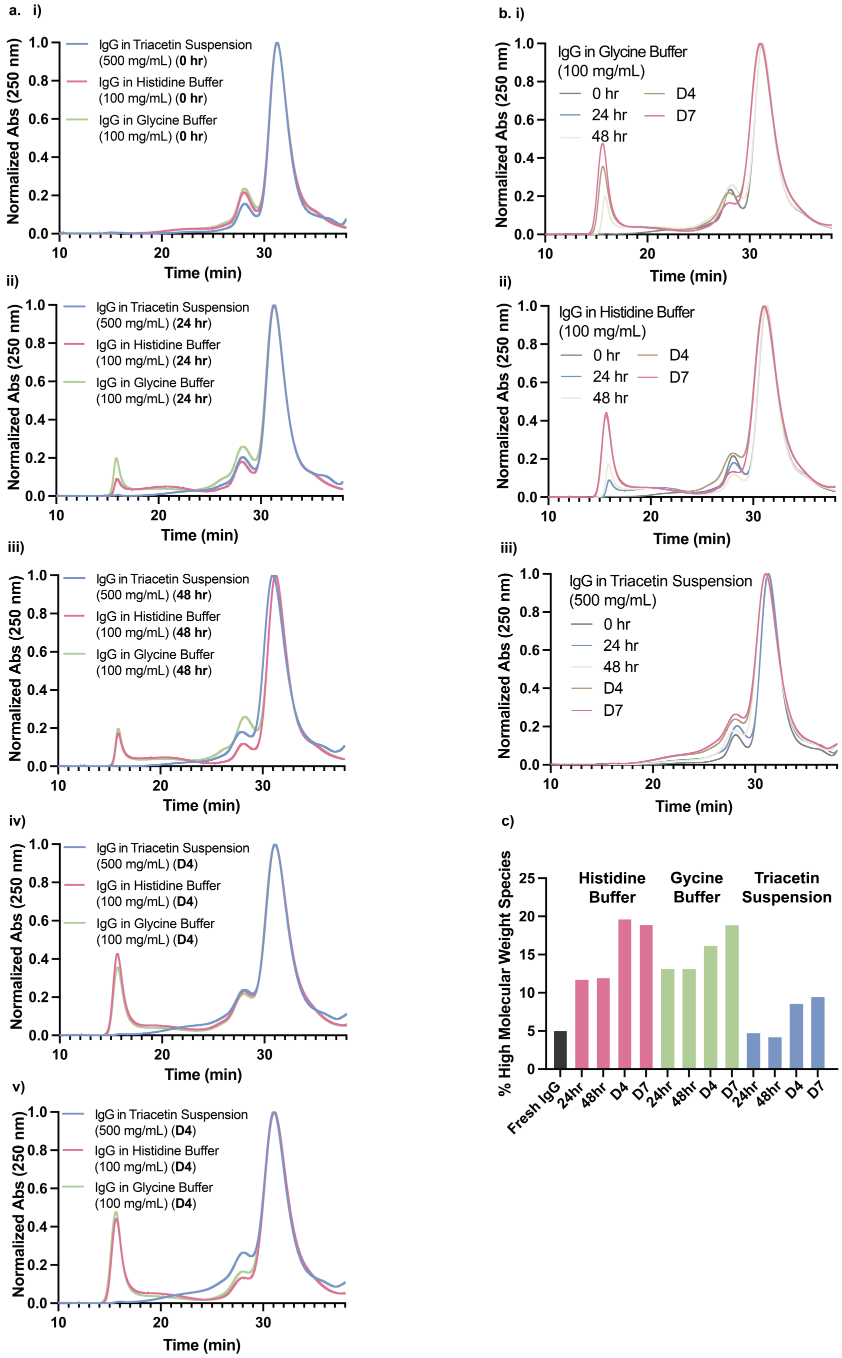

**SI Figure 10. Stressed aging of IgG. a)** SEC of hIgG in triacetin (500 mg/mL), hIgG in histidine buffer (100 mg/mL) and hIgG in glycine buffer (100 mg/mL) following **i)** 0 hr, **ii)** 24 hr, **iii)** 48 hr, **iv)** 4 days, and **v)** 7 days of stressed aging at 50 °C with shaking. **b)** Comparative SEC over 7 days of stressed aging for **i)** hIgG in glycine buffer (100 mg/mL), **ii)** hIgG in histidine buffer (100 mg/mL), and **iii)** hIgG in triacetin (500 mg/mL) and **c)** corresponding high molecular weight fraction for each formulation following stressed aging.

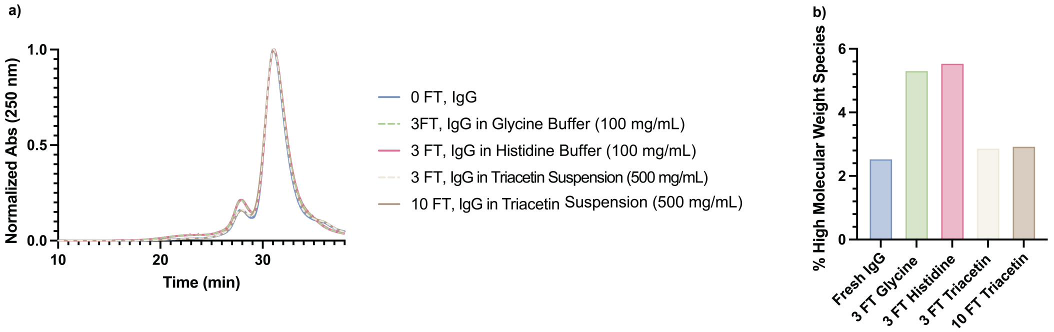

**SI Figure 11. IgG stability through repeated freeze-thaw (FT) cycles. a)** SEC of hIgG in triacetin (500 mg/mL), hIgG in histidine buffer (100 mg/mL) and hIgG in glycine buffer (100 mg/mL) following repeated freeze thaw cycles and **c)** corresponding high molecular weight fraction for each formulation.

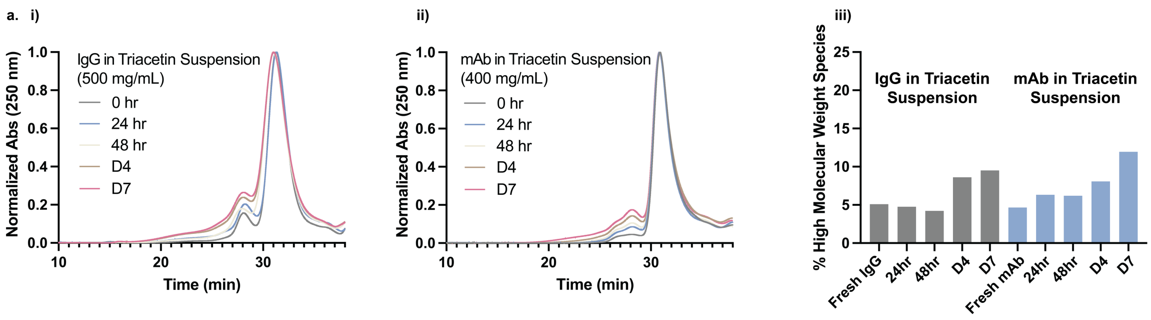

**SI Figure 12. Comparative stressed aging of hIgG and mAb triacetin suspensions. a)** SEC of **i)** hIgG in triacetin (500 mg/mL) and **ii)** human mAb in triacetin (400 mg/mL) and **iii)** corresponding high molecular weight fraction for each formulation.

**a) Subcutanous injectable liquid high concentration antibody products approved in the US between 1998 - October 2021**

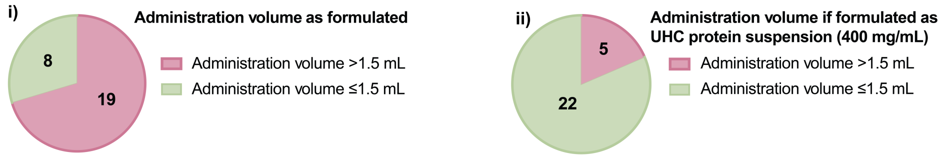

**b) Comercialized high concentration antibody products where formulation as UHC protein suspensions would enable dosing in a single 1.5 mL injection**

| Therapeutic Protein | Brand Name | Therapeutic Dose (mg) | Concentration (mg/mL) | Injection Volume as Formulated (mL) | Required Needle Pricks as Formulated | Injection Volume if Formulated as UHC Suspension (400 mg/mL) (mL) | Required Needle Pricks as UHC Suspension |
|---|---|---|---|---|---|---|---|
| omalizumab | Xolair® | 600 | 150 | 4 | 3 | 1.5 | 1 |
| dupilumab | Dupixent® | 600 | 150 | 4 | 3 | 1.5 | 1 |
| trastuzumab | Herceptin Hylecta® | 600 | 120 | 5 | 4 | 1.5 | 1 |
| certolizumab pegol | Cimzia® | 400 | 200 | 2 | 2 | 1 | 1 |
| canakinumab | Ilaris® | 300 | 150 | 2 | 2 | 0.75 | 1 |
| golimumab | Simponi® | 200 | 100 | 2 | 2 | 0.5 | 1 |
| evolocumab | Repatha® | 420 | 140 | 3 | 2 | 1.05 | 1 |
| galcanezumab-gnlm | Emgality® | 240 | 120 | 2 | 2 | 0.6 | 1 |
| lanadelumab-flyo | Takhzyro® | 300 | 150 | 2 | 2 | 0.75 | 1 |
| Adalimumab | Humira® | 160 | 100 | 1.6 | 2 | 0.4 | 1 |
| belimumab | Benlysta® | 400 | 200 | 2 | 2 | 1 | 1 |
| emicizumab-kxwh | Hemlibra® | 400 | 150 | 2.67 | 2 | 1 | 1 |
| alirocumab | Praluent® | 300 | 150 | 2 | 2 | 0.75 | 1 |
| secukinumab | Cosentyx® | 300 | 150 | 2 | 2 | 0.75 | 1 |

**SI Figure 13. Potential to reduce injection burden for commercial HCA products.** Solution-based high-concentration antibody (HCA) products (>100 mg/mL) approved by the FDA between 1998 and 2021 that can be administered by subcutaneous injection. **a. i)** Fraction of HCA products as formulate where a therapeutic dose can be administered with a volume of 1.5mL. **ii)** Fraction of HCA products formulated as 400 mg/mL UHC suspensions where a therapeutic dose can be administered with a volume of 1.5mL. **b)** Commercialized HCA products where formulation as a UHC protein suspension would enable dosing in a single 1.5mL injection.

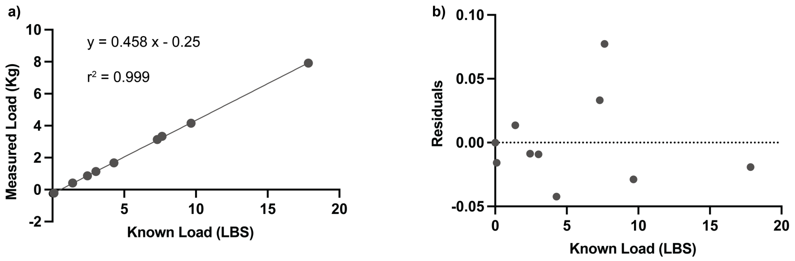

**SI Figure 14. Calibration of FUTEK force sensor. a)** Linear relationship between known load and force sensor measured load. Injection force measurements typically fall between 0 and 4 Kg. To confirm the calibration of the FUTEK force sensor, objects of a known weight in pounds were placed onto the force sensor, and weights were measured in Kg. The slope of the line (m=0.45, $R^2$ = 0.99) agrees with the conversion of 1 lb = 0.45 kg. The slight shift in y intercept is accounted for during data analysis by subtracting the baseline injection force from the recorded plateau injection force. **b)** Residuals of best fit line.

### Supplementary Discussion

#### *Flow behavior of suspensions*

The flow behavior of high solid fraction suspensions can be described with the Krieger & Dougherty model (1959). This model includes the terms suspension viscosity (η), non-solvent viscosity ($\eta_0$), intrinsic viscosity ([η]), particle volume fraction (Φ), and max particle volume fraction ($\Phi_{max}$). Intrinsic viscosity is a measure of particle shape. It is fixed at 2.5 for spherical particles but can be greater than 2.5 for particles with increased surface irregularities. Because injection force is proportional to viscosity, we fit our injection force data to a modified version of the Krieger & Dougherty model where $F_{inj} \propto \eta$. Here, the term for liquid viscosity ($\eta_0$) is instead fixed as the injection force of the pure non-solvent (70:30 Triacetin: DMAc, or 100 Triacetin) ($F_{inj}0$). Injection force data as a function of particle volume fraction is fit to the Krieger & Dougherty model (1959) to solve for $\Phi_{max}$ and [η]. For spherical particles, $\Phi_{max}$ ranges between 0.52 and 0.74, and higher $\Phi_{max}$ values indicate the suspension is closer to achieving maximum particle loading. $\Phi_{max}$ can exceed 0.74 when particles have increased polydispersity as the packing space can be filled more efficiently.

#### *Krieger & Dougherty model:*

$$\eta = \eta 0 \left(1 - \frac{\Phi}{\Phi \mathrm{max}}\right) - [\eta] * \Phi \mathrm{max}$$

Suspension viscosity: $\eta$

Non-solvent viscosity: $\eta 0$

Intrinsic viscosity: [η]

Particle volume fraction: $\Phi$

Max particle volume fraction: $\Phi\mathrm{max}$

#### *Assumptions for Modified Krieger & Dougherty model:*

1. Injection force is proportional to viscosity. $F_{inj} \propto \eta$
2. If injection force is proportional to viscosity, then the injection force of non-solvent without particles is proportional to the non-solvent viscosity. $F_{inj}0 \propto \eta 0$

#### *Modified Krieger & Dougherty model:*

$$F_{inj} = F_{inj}0 \left(1 - \frac{\Phi}{\Phi \mathrm{max}}\right) - [\eta] * \Phi \mathrm{max}$$

Injection force: $F_{inj}$

Non-solvent injection force: $F_{inj}0$

Intrinsic viscosity: [η]

Particle volume fraction: $\Phi$

Max particle volume fraction: $\Phi\mathrm{max}$

#### *Data fitting:*

Experimental data shows injection force ($F_{inj}$) as a function of particle volume fraction ($\Phi$). Non-solvent injection force ($F_{inj}0$) is fixed as the experimental injection force value for pure non-solvent. Intrinsic viscosity ([η]) is initially fixed at 2.5 for spherical particles and allowed to vary as necessary. The data is fit to the modified Krieger & Dougherty model and fitting results in a value for maximum particle volume fraction ($\Phi_{max}$).
